# Evidence of centromeric histone 3 chaperone involved in DNA damage repair pathway

**DOI:** 10.1101/2024.07.26.605363

**Authors:** Prakhar Agarwal, Santanu Kumar Ghosh

## Abstract

The centromeric protein-A (CENP-A) is an evolutionary conserved histone H3 variant that marks the identity of the centromeres. Several mechanisms regulate the centromeric deposition of CENP-A as its mislocalization causes erroneous chromosome segregation, leading to aneuploidy-based diseases, including cancers. The most crucial deposition factor is a CENP-A specific chaperone, HJURP (Scm3 in budding yeast), which specifically binds to CENP-A. However, the discovery of HJURP as a DDR (DNA damage repair) protein and evidence of its binding to Holliday junctions in vitro indicate a CENP-A-deposition-independent role of these chaperones. In this study, using budding yeast, we demonstrate that Scm3 is crucial for the DDR pathway as *scm3* cells are sensitive to DNA damage. We further observe that the *scm3* mutant interacts with the *rad52* DDR mutant and is compromised in activating DDR-mediated arrest. We demonstrate that Scm3 associates with the DNA damage sites and undergoes posttranslational modifications upon DNA damage. Overall, from this report and earlier studies on HJURP, we conclude that DDR functions of CENP-A chaperones are conserved across eukaryotes. Thus, the revelation that these chaperones confer genome stability in more than one pathway has clinical significance.

## Introduction

The faithful replication and partitioning of the replicated chromatin as chromosomal units from mother to daughter cell is important for cell survival. New nucleosomes, the units of chromatin, are assembled on DNA during replication that requires deposition of the histones within the nucleosomes in a highly regulated step-by-step process that involves the deposition of (H3-H4)_2_ tetramer followed by two H2A-H2B dimers (Hammond et al., 2017; Ransom et al., 2010). The deposition of the histones to their specific positions within the nucleosome occurs in a temporal manner assisted by a group of proteins called histone chaperones. For this, the chaperones physically interact with the histones in a pre-nucleosomal complex and thus prevent premature histone-DNA interaction or nucleosome assembly (Burgess and Zhang, 2013; Ransom et al., 2010). The deposition of the canonical histones at the gross chromatin takes place mostly during DNA replication via a replication-coupled (RC) pathway using the chaperones dedicated to the RC pathway (Serra-Cardona & Zhang, 2018). For instance, Asf1 and CAF-1 deposit (H3-H4)_2_ tetramer (Burgess and Zhang, 2013; De Koning et al., 2007) and Nap1 facilitates (H2A-H2B)_2_ assembly (De Koning et al., 2007; Eitoku et al., 2008) into the nucleosomes. On the other hand, the variant histones that earmark specialized chromatin domains committed to do specific biological functions are deposited at specific sites throughout the cell cycle using replication-independent (RI) pathway using variant-specific chaperones dedicated to RI pathway (Smith, 2002). For Instance, deposition of the variant histones H2A.Z, centromeric H3 (CenH3), MacroH2A signifying transcriptional activation (Giaimo et al., 2019), kinetochore formation (Earnshaw and Rothfield, 1985; Earnshaw and Tomkiel, 1992; McKinley and Cheeseman, 2016) and X chromosome inactivation (Sun and Bernstein, 2019) occurs via chromatin remodeler (Fan et al., 2022), HJURP chaperone (Dunleavy et al., 2009; Foltz et al., 2009) and acidic nuclear phosphoprotein 32B (ANP32B) (Mandemaker et al., 2023), respectively.

Almost all the eukaryotes harbor CenH3 at the centromeres, which promotes the formation of kinetochore at those loci (Blower and Karpen, 2001; Buchwitz et al., 1999; Howman et al., 2000; Stoler et al., 1995; Van Hooser et al., 2001). Since the kinetochores bind to the microtubules and drive faithful chromosome segregation, deposition of CenH3 at the centromeres is essential for cell survival. In metazoans, CENP-A (Centromeric protein-A) is the CenH3 (Earnshaw and Rothfield, 1985), and HJURP (Holliday junction recognition protein) is the cognate chaperone (Dunleavy et al., 2009; Foltz et al., 2006). The CENP-A binding domain within HJURP present at its N-terminal is highly conserved (Aravind et al., 2007; Sanchez-Pulido et al., 2009; Shuaib et al., 2010). Despite the well-established function of HJURP in CENP-A deposition at the centromeres, HJURP was initially identified as a protein involved in DNA damage response (DDR) due to the following reasons (Kato et al., 2007). HJURP was so named because of its ability to bind in vitro to Holliday junction DNA substrates that are in vivo generated during DNA damage repair through homologous recombination. In addition, HJURP was found to interact physically with DNA damage sensing complex MRN (Mre11-Rad50-Nbs1), and the interaction was found to increase significantly upon induction of double-strand breaks (DSBs). As a corollary, the level and subcellular localization of HJURP change in the cells treated with DNA-damaging agents such as Infrared Radiation (IR), Hydroxyurea (HU), cisplatin, and camptothecin (CPT). Finally, the expression of HJURP was found to be regulated by ATM, a DDR kinase, as no expression was observed in ATM-deficient cancerous cell lines. Perhaps due to genome wide function on DDR, HJURP has been visualized to be diffusely present throughout the nucleus (Dunleavy et al., 2009; Kato et al., 2007).

Given the functions of HJURP in CENP-A deposition and DDR pathway, it is not surprising that misregulation of HJURP is involved in disease states. HJURP is considered an oncogene as its overexpression has been associated with several cancers (Chen et al., 2018; Kang et al., 2020) including breast cancer (Coates et al., 2010; Hu et al., 2010; Montes de Oca et al., 2015), lung cancer (Kato et al., 2007), ovarian cancer (Dou et al., 2022), glioblastoma (Valente et al., 2013, 2009), hepatocellular carcinoma (Chen et al., 2018; Hu et al., 2017; Luo et al., 2022), colorectal cancer (Kang et al., 2020) and in pancreatic cancer (Wang et al., 2020). Consequently, the inhibition of HJURP attenuates the properties of the cancer cells and increases the survival rate of the patients, and thus, HJURP acts as a prognostic marker for several cancers (Kang et al., 2020; Lai et al., 2021).

In budding yeast, Cse4 and Scm3 are the homologs of CENP-A and HJURP, respectively. Scm3 deposits Cse4 at the centromeres during the S phase through its physical interaction with Ndc10 that binds to the centromeric DNA as a part of the CBF3 complex (Camahort et al., 2007; Cho and Harrison, 2012, 2011). It has been shown that following the deposition of Cse4, Scm3 remains associated with the centromeres throughout the cell cycle (Wisniewski et al., 2014; Xiao et al., 2011). Like HJURP, Scm3 has been found on the gross chromatin besides its localization at the centromeres (Mizuguchi et al., 2007). Furthermore, a comparison of the domain organization between HJURP and Scm3 suggests that besides the highly conserved CENP-A binding domain, both HJURP and Scm3 also harbour homologous DNA binding domain (Figure S1), which is predicted to be evolutionarily conserved in all vertebrates and fission yeast (Müller et al., 2014). Notably, overexpressed Scm3, like HJURP, shows chromosome instability, and it is believed that Scm3, when excess, binds to the centromeric DNA independently and leads to chromosome missegregation (Choy et al., 2012; Mishra et al., 2011). Both the proteins dimerizes and in *in-vitro* assay, they can bind even to non-*CEN* DNA (Kato et al., 2007; Xiao et al., 2011). While there is no mammalian homolog of budding yeast Ndc10 and no yeast homolog of mammalian Mis18, both these proteins are critical for CENP-A deposition as they bind to their respective CENP-A chaperones through regions independent of CENP-A or DNA binding domain. Therefore, despite the large difference in the polypeptide length between HJURP and Scm3 (748 vs. 229), most of the functions are highly conserved from humans to yeast. Given the physical interaction of HJURP with the MRN complex, it is highly plausible that Scm3 might also interact with DDR proteins in yeast. Altogether, from the above observations, we hypothesize that Scm3, like HJURP, may also be involved in the DDR pathway in budding yeast. In this work, we report that the cells lacking Scm3 are sensitive to the DNA damaging agents and are compromised both in DNA damage repair. In addition to the known functions of HJURP in DDR pathway, in this report we show, using Scm3, that these chaperones are required for mounting DDR-dependent checkpoint activation. Importantly, we demonstrate for the first time that they indeed associate with the DNA damage sites upon induction of DNA damage and persists there perhaps to facilitate damage repair. Our results suggest that the phosphorylation of Scm3 may be attributed to the DDR functions of Scm3. Taken together, this study reveals an evolutionary kinship amongst CenH3 chaperones in conferring genome stability through efficient DNA damage repair, besides their canonical role in CenH3 deposition.

## Results

### Scm3 depleted cells are sensitive to MMS mediated DNA damage

The yeast cells compromised in DNA damage repair (DDR) function show sensitivity to DNA damaging agents such as methyl methane sulfonate (MMS), HU and CPT. To investigate the role of Scm3 in DDR, we constructed an auxin-inducible (Nishimura et al., 2009) Scm3 conditional mutant since the protein is essential for growth (Camahort et al., 2007; Mizuguchi et al., 2007; Stoler et al., 2007), and assayed the behaviour of the mutant in the presence of the DNA damaging agents. Since the cells devoid of the homologous recombination protein, Rad52, are known to show extreme sensitivity to such drugs, *rad52Δ* cells were taken as a positive control in our assays (Hallwyl et al., 2008; Kaytor and Livingston, 1994). Mid-log grown cells were harvested, serially diluted, and spotted on plates supplemented with auxin and DNA damaging agents (MMS, HU, CPT) in different combinations and found similar results for all the drugs (Figure 1A, Figure S2A). We observed that *scm3* mutants (three independent transformants) were more sensitive in the presence of MMS (plate supplemented with both auxin and MMS) as compared to those without MMS (plate supplemented with only auxin), whereas *rad52Δ* cells, as expected, were sensitive to MMS but not to auxin (Figure 1A). To quantify the levels of sensitivity towards MMS, we performed a CFU count assay where *SCM3-AID* cells were pregrown in rich media without auxin till the mid-log phase before an equal number of cells were plated on media supplemented with auxin and MMS in different combinations as indicated in Figure 1A. The number of colonies formed after 36 hrs were counted and are quantified as follows. The number of colonies that appeared on the plates free of auxin and MMS were taken as 100% cell viability; with respect to this, the number of colonies that appeared on other plates (conditions) was used to calculate the viability percentage for those conditions. We observed a statistically significant drop in cell viability in the plate supplemented with both auxin and MMS compared to auxin alone (Figure 1B).

**Figure 1.**
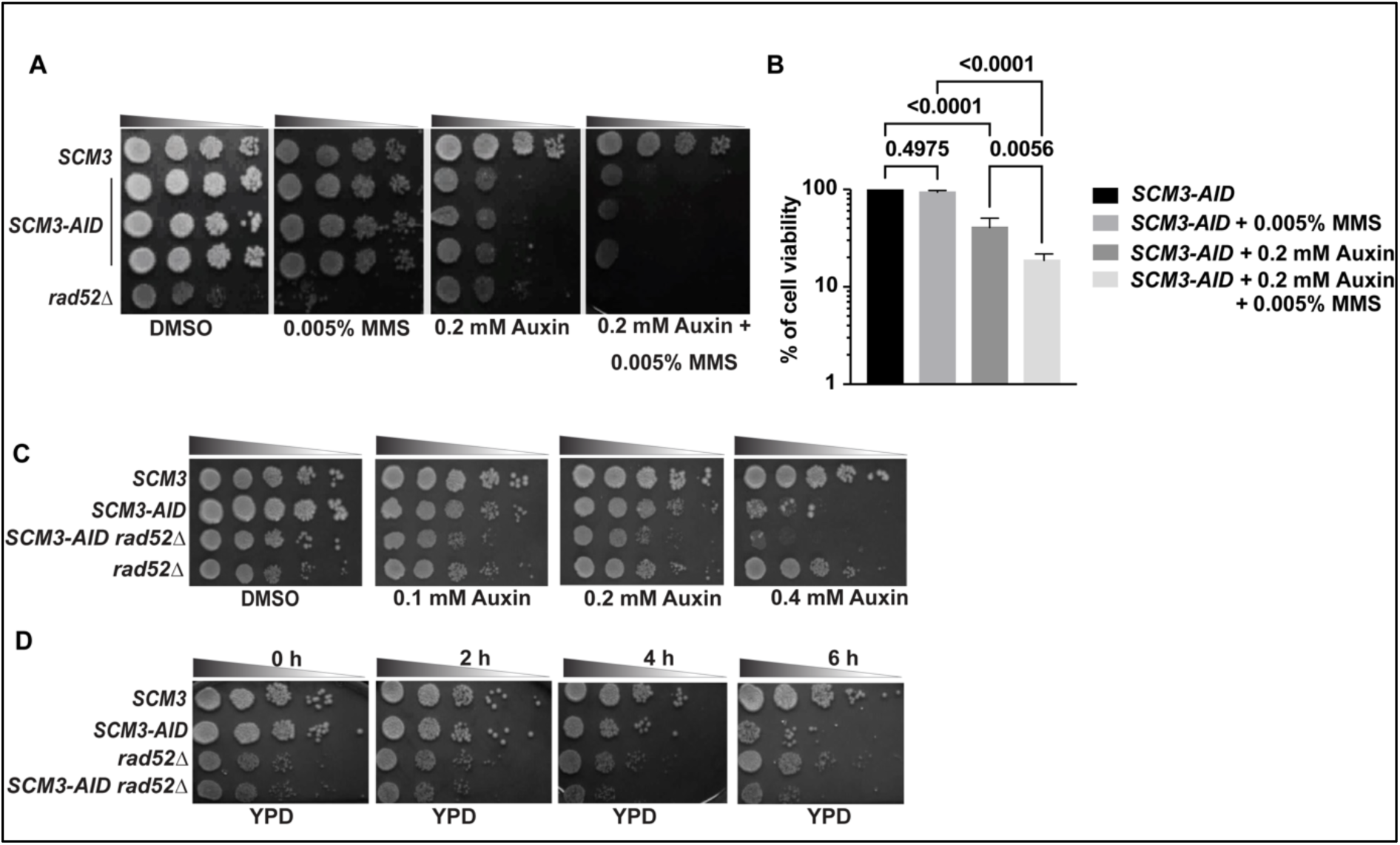
Loss of Scm3 causes sensitivity to MMS-mediated DNA damage. **(A)** Wild-type (*SCM3*), *SCM3-AID* (three independent transformants)*, and rad52Δ* cells grown till mid-log were serially diluted and spotted on the indicated plates. The plates were incubated at 30°C for 24-48 hrs before imaging. **(B)** An equal number of mid-log grown *SCM3-AID* cells were spread on plates containing indicated supplements. From the number of colonies that appeared after incubation at 30°C for 2 days, the cell viability was estimated (see materials and methods) and plotted. Colonies formed on the plate without auxin and MMS were taken as 100% viability, and accordingly, viability percentage was calculated for other plates. The *p* values were estimated by one-way ANOVA test from three independent experiments. **(C)** Wild-type (*SCM3)*, *SCM3-AID, SCM3-AID rad52Δ,* and *rad52Δ* cells grown till mid-log were serially diluted and spotted in the presence of indicated auxin concentrations. The plates were incubated at 30°C for 24-48 hrs before imaging. **(D)** The cells of the strains used in (C) were grown till mid-log and treated with 0.75 mM auxin for the indicated duration before they were washed, serially diluted, and spotted on YPD plates that were incubated at 30°C for 24-48 hrs before imaging.

Since the presence of auxin in the plate can hinder the growth of the *SCM3-AID* cells, in another approach to examine the sensitivity of *scm3* mutant to MMS, we pre-treated the cells with 0.75 mM auxin for 2 hrs to deplete Scm3 before they were washed and spotted on plates containing different concentrations of MMS, but without auxin. The depletion of Scm3 was verified by observing higher percentage of G2/M arrested cells (Figure S2B). Here also we found that Scm3-depleted cells were more sensitive to MMS as compared to the wild-type cells (Figure S2C). To quantify the sensitivity observed by this approach, we performed a CFU count assay as above. Following auxin pretreatment, an equal number of wild-type, *SCM3-AID,* and *rad52Δ* cells were spread on the YPD plate or YPD plate containing 0.01% MMS. By calculating the viability percentage as described above, we again found a statistically significant drop in cell viability on the plate where Scm3 was predepleted (Figure S2D). The *rad52Δ* cells, as the positive control, showed high sensitivity, as expected.

The increased sensitivity of *scm3*-depleted cells to DNA-damaging agents could be due to weakening of the kinetochores as Scm3-mediated deposition of Cse4 promotes kinetochore assembly (Camahort et al., 2007; Cho and Harrison, 2011). If this holds true, perturbation of the kinetochore by degradation of other kinetochore proteins must show a similar sensitivity to MMS. In budding yeast, Ndc10 is recruited to the centromeres upstream of Scm3 (Lang et al., 2018), whereas the centromeric localization of Mif2, another essential inner kinetochore protein, depends on Scm3 and Cse4 (Xiao et al., 2017). We constructed *NDC10-AID* and *MIF2-AID* strains and used them for our assay to represent the proteins independent or dependent on Scm3 for centromeric localization, respectively. We also included one non-essential kinetochore protein, Ctf19, a protein of the COMA complex, to remove any possible mis-judgement in distinguishing cell-growth-arrest phenotype occurring due to drug-sensitivity vs. auxin-mediated degradation of essential proteins. The COMA complex is directly recruited to the centromeres through interaction with the N terminal tail of Cse4, hence dependent on Scm3 (Chen et al., 2000; Fischböck-Halwachs et al., 2019). Mid-log grown cells were harvested and spotted on the indicated plates, however, we did not observe any increased sensitivity of such cells to MMS (Figure S3). In fact, Mif2 depleted cells showed a slight resistance to MMS due to unknown reasons. Therefore, the increased sensitivity to MMS in *scm3* mutant but not in other kinetochore mutant cells indicates that Scm3 possesses an additional function in genome stability besides its role in kinetochore assembly.

Since HJURP is involved in DDR through the HR pathway (Kato et al., 2007) and If Scm3 mimics HJURP to repair damaged DNA, then the *scm3* mutant is expected to interact genetically with the HR mutants. We tested that possibility by analyzing the genetic interaction between *SCM3* and *RAD52,* as Rad52 is essential for efficient homologous recombination in *S. cerevisiae* (Barlow & Rothstein, 2010; Lee et al., 2003). We deleted *RAD52* in the *SCM3-AID* strain, and the mid-log grown cells were spotted on plates supplemented with increasing concentrations of auxin (Figure 1C). Interestingly, the double mutant grew slowly as compared to the single mutants and the wild-type strain. To further validate the genetic interaction between the two genes, the mid-log grown cells were pre-treated with a 0.75 mM auxin to deplete Scm3 for the indicated duration before the cells were spotted on YPD plates. Again, we observed that the double mutant grew slower than the wild-type and single mutants (Figure 1D). Next, we wished to investigate how the double mutant behaves when challenged with DNA damaging agent. As expected, when the double mutant was spotted in the presence of 0.005% MMS, all the single mutants grew better than the double mutant (Figure S2E). These results indicate a genetic interaction between *SCM3* and *RAD52,* which becomes pronounced in the presence of DNA-damaging agents.

Further, we wanted to identify the regions in Scm3 whose loss confers sensitivity to DNA damage. The domain organization of Scm3 reveals two essential conserved regions, namely NES (nuclear export signal) and HR (heptad repeat), having functions in DNA and Cse4 binding, respectively (Figure S1A). Scm3 has a non-essential D/E-rich region, which is important for protein stability but not involved in Cse4 binding (Stoler et al., 2007). Scm3 also possesses two BR (bromodomain) regions, also referred to as the NLS (nuclear localization signal). The deletion of the C-terminal 25 amino acids or the bromodomain regions does not cause cell lethality (Shivaraju et al., 2011; Stoler et al., 2007). We used the versions of Scm3 mutants used elsewhere (Shivaraju et al., 2011) harbouring deletion in one or the other non-essential region as the sole source of Scm3 and challenged these mutants (two independent transformants) to MMS and CPT along with wild-type Scm3 as a control. We found no growth defect in response to MMS or CPT (Figure S1B) suggesting that at least the nonessential regions are not important for the DDR function of Scm3. Although this indicates that the essential NES and/or HR might be involved but that is technically difficult to test using growth based assays.

From these results, we conclude that the loss of Scm3 confers sensitivity to DNA damaging agents, indicating that Scm3, much like its mammalian homolog HJURP, may play a role in the DDR pathway.

### The loss of Scm3 generates more Rad52 foci

Having shown that the *scm3* mutant is sensitive to DNA damage, we hypothesized that the mutant cells might be deficient in repairing intrinsic and extrinsic DNA damages. To examine this, we wished to visualize the Rad52-GFP foci in these cells with or without MMS-induced DNA damage. Since in budding yeast, the primary pathway to repair DNA damage is through homologous recombination (HR) mediated by Rad52, deficiency in repair will cause accumulation of DNA lesions and hence Rad52 (Dornfeld and Livingston, 1991; Lisby et al., 2003, 2001). The mid-log grown *RAD52-GFP SCM3-AID* cells were split into auxin- and DMSO-treated halves. Following 2 hrs of treatment, the cells were again split and were mock-treated or treated with 0.02% MMS for 90 mins (Figure 2A). We visualized the formation of Rad52-GFP foci in these cells (Figure 2B). Interestingly, we observed that 18% of the Scm3 depleted (+auxin) cells showed Rad52-GFP foci in contrast to only 5% in the wild type (Figure 2C, untreated). In accord with this, when these cells were challenged with MMS, a significant increase of cells with Rad52-GFP foci was observed in the mutant (70%) compared to the wild type (45%) (Figure 2C, 0.02% MMS). Moreover, while the mutant cells harbored an average of ∼ 3 foci, the wild type showed ∼2 foci per cell (Figure 2D) in response to MMS treatment. These results suggest that both intrinsically and extrinsically generated DNA lesions remain unrepaired in the absence of Scm3.

**Figure 2.**
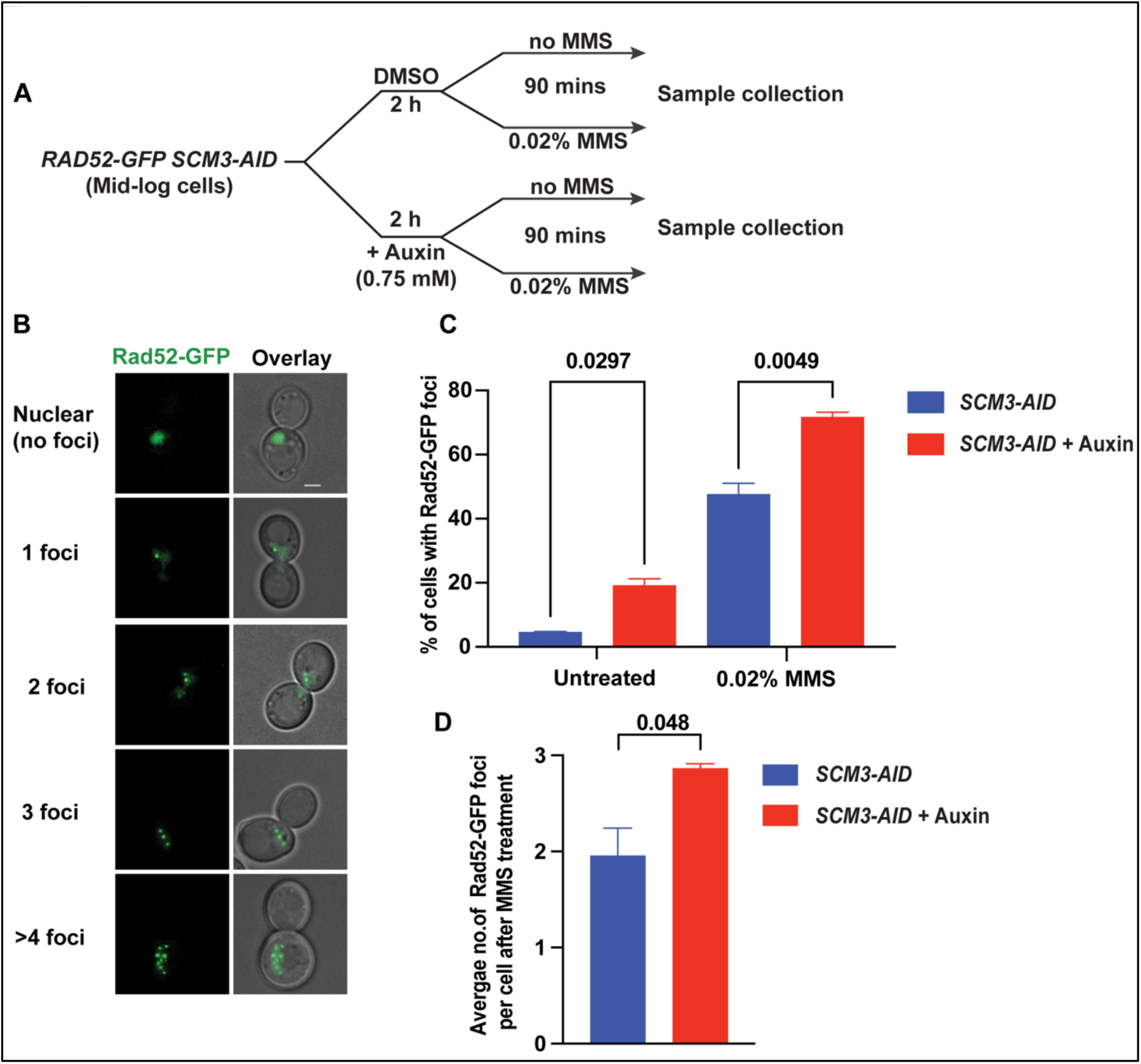
Loss of Scm3 causes increased Rad52 foci in the presence or absence of MMS. **(A)** The schematic of the experimental strategy that was followed for visualization of Rad52-GFP foci in the *RAD52-GFP SCM3-AID* cells either in the presence of DMSO (wild-type) or auxin (Scm3 depleted) and each either treated with MMS or left untreated. **(B)** Representative images showing different patterns of Rad52-GFP foci. Rad52-GFP produces a nuclear signal (row 1) in the absence of any DNA damage, but with the damage, it coalesces to form bright foci (rows 2-5) representing the DNA repair sites. **(C)** The percentage of indicated cells containing Rad52-GFP foci with or without MMS treatment for 90 mins. **(D)** The average number of Rad52-GFP foci per MMS-treated cell from the experiment described in (C). At least 300 cells were analyzed from three independent experiments for each set in **(C)** and **(D)**. The *p* values were estimated by unpaired two-tailed student’s t-test. Scale bar = 2 µm.

To understand the dynamics of Rad52 foci formation in *scm3* cells, we used a strategy as shown in Figure S4A. Briefly, the *bar1Δ RAD52-GFP SCM3-AID* cells were arrested in the G1 phase of the cell cycle by adding alpha factor. 1 hour before release from the G1 phase, cells were treated with auxin to deplete Scm3 or with DMSO control. Subsequently, the cells were washed several times and released into alpha factor free media supplemented with or without MMS but in the presence of nocodazole in order to have all the cells eventually arrested at metaphase. Following release, the cells were harvested every 30 mins and the percentage of cells containing Rad52-GFP foci were visualized (Figure S4B). Scm3 depleted cells, but not the wild type, with or without MMS, showed a peak of Rad52-GFP accumulation at around 60 mins from G1 release (Figure S4C, D). Notably, this time point possibly coincides with the timing of DNA replication (judged by bud morphology) when DNA is most susceptible to intrinsic and extrinsic damages. Taken together, Scm3-depleted cells exhibit more Rad52 foci, indicating a compromised DDR pathway in these cells.

If the DNA lesions remain unrepaired, the cells might undergo an apoptotic pathway leading to cell death. To determine the percentage of cell death in the *scm3* mutant culture, the cells harvested at 90 mins of MMS or mock treatment (after auxin treatment), as shown in Figure 2A were stained with propidium iodide (PI), which does not permeate the live cells and stains only the dead cells (Figure S4E, arrow). We observed that while ∼15% of the Scm3 depleted (+auxin, -MMS) cells were PI positive, it was only ∼4% for the wild type (DMSO, - MMS) (Figure S4E, untreated). The PI positive cell population increased after MMS treatment in both the cultures with significant difference between them (35% vs 27%) (Figure S4E, 0.02% MMS). These results suggest that unrepaired intrinsic and extrinsic DNA lesions in the absence of *scm3* lead to compromised cell viability.

### Scm3 associates with non-centromeric sites in response to DNA damage

Having observed that Scm3 depleted cells are compromised in DNA damage repair and *scm3* interacts genetically with *rad521*, it is reasonable to argue that Scm3 goes to the DNA damage sites to promote repair. To examine this, we first wanted to see if Scm3 alters its localization pattern in response to MMS-mediated DNA damage. Since Scm3 is also expected to localize at the kinetochores (Mizuguchi et al., 2007), we visualized Scm3 along with the kinetochore marker, Ndc10, on chromatin spreads. The cells harboring *NDC10-6HA SCM3-13MYC* were mock-treated or treated with 0.02% MMS for 90 mins. Ndc10 showed bright foci representing the kinetochore cluster, whereas Scm3 was localized to multiple locations across chromatin, including its colocalization with Ndc10 (Figure 3A, arrow). Similar localization of Scm3 has been observed earlier in live cells where Scm3-GFP showed a wider nuclear localization, which might argue for its genome-wide functions (Luconi et al., 2011; Wisniewski et al., 2014). We also observed a significant enrichment of Scm3 at the rDNA loops irrespective of the extrinsic DNA damage (Figure 3A, arrowhead). The enrichment of Scm3 at the rDNA loops is perhaps to maintain the highly repetitive organization of rDNA. HJURP has also been shown to localize rDNA arrays in mammalian systems (Kato et al., 2007). Additionally, upon quantifying the total intensity of Scm3-13Myc and Ndc10-6HA and normalizing that with the background intensity, we observed a statistically significant increase in the Scm3-13Myc, but not Ndc10-6HA intensity in response to MMS treatment (Figure 3B). No change in the intensity of Ndc10-6HA post-MMS treatment justifies the absence of increased sensitivity of Ndc10 depleted cells in response to MMS (Figure S3A).

**Figure 3.**
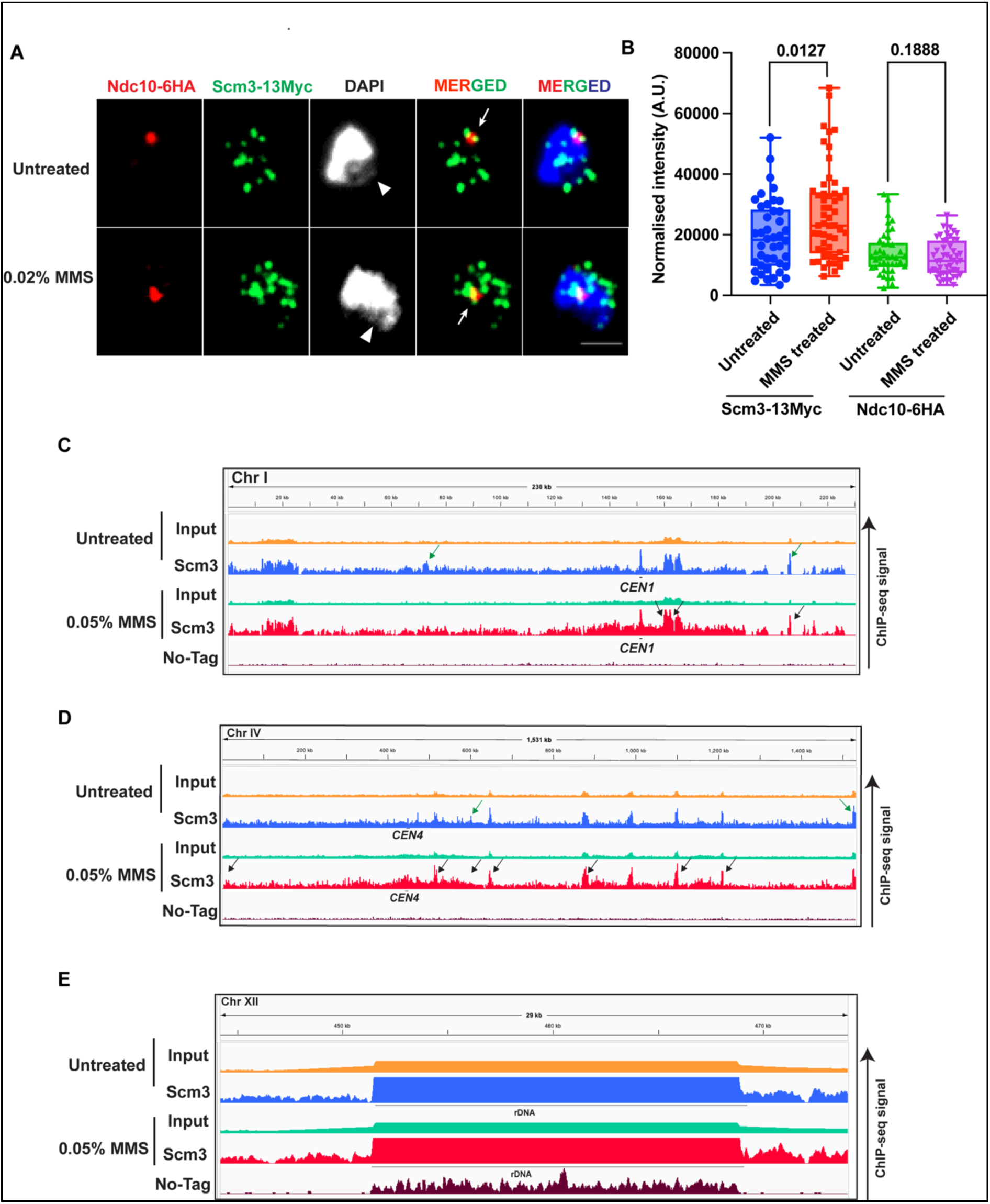
The localization of Scm3 in response to DNA damage. **(A)** Representative images showing the localizations of Ndc10-6HA and Scm3-13Myc using chromatin spread. The mid-log grown *NDC10-6HA SCM3-13MYC* cells were treated with 0.02% MMS or mock-treated for 90 mins for the assay. The arrow shows the co-localization of Ndc10-6HA and Scm3-13Myc at the kinetochore cluster, and the arrowhead shows the rDNA loop. **(B)** The intensities of Scm3-13Myc and Ndc10-6HA normalized with the background intensity were quantified using ImageJ software and were plotted. At least 30 spreads from 2 independent experiments were analyzed for each set in **(B).** The *p* values were estimated by unpaired two-tailed student’s t-test. Scale bar = 2 µm. **(C, D)** The ChIP-seq signal of Scm3 (Scm3-13Myc) and input from cells either untreated or treated with 0.05% MMS for the entire chromosome I; the no-tag strain was used as a control. The plot shows the binding of Scm3 across (C) chromosome I and (D) chromosome IV. The green and black arrows represent the non-centromeric localization of Scm3 in untreated, and MMS treated samples, respectively. **(E)** Same as (C); only a portion of chromosome XII flanking 9.1 kb rDNA region is shown. The scale is 7–50 for the input samples and 7-25 for IP samples.

Having observed an increased chromatin localization of Scm3 in response to MMS on chromatin spreads, we wished to verify this by determining genome wide association of Scm3 using ChIP-Seq. To negate any effect on Scm3’s binding due to cell cycle progression we first arrested the cells harboring *SCM3-13MYC* at metaphase by nocodazole. The arrested cells were further mock treated or treated with 0.05% MMS for an additional 90 mins before they were harvested for ChIP. The Scm3 ChIP-seq signal was normalized with the input signal, and genome-wide plots were generated to see Scm3 binding along the entire chromosomes. We also performed ChIP-seq from a no-tag control strain that provided the background signal due to the antibody or the beads treatment. We could observe an enrichment of Scm3 in both MMS treated and untreated cells at the centromeres, while no enrichment in the no-tag sample was observed (Figures 3C, 3D, S5). In the untreated sample, the peaks called by MACS2 after input normalization showed enrichment of Scm3 at a few other genomic locations as well besides the centromeres (Figures 3C, 3D, S5, green arrows), which was also observed in our chromatin spread data (Figures 3A, 3B). Interestingly, in MMS treated cells, following same methodology, we could observe a greater number of peaks of Scm3 binding to the non-centromeric locations (Figures 3C, 3D, S5, black arrows). While in the untreated sample, Scm3 localized to 1-2 non-centromeric locations, the same increased drastically in the treated sample. Such non-centromeric chromatin enrichment of Scm3 was observed for several chromosomes (S5, black arrows) and corresponds to the increased chromatin localization of Scm3 in response to MMS, as visualized on chromatin spreads (Figure 3B). Further, we could observe the localization of Scm3 at the 9.1 kb rDNA region on ChrXII, validating our chromatin spread data (Figure 3E).

### Scm3 partially co-localizes with Rad52, perhaps at the DNA damage sites

Given the increased chromatin association of Scm3 in response to DNA damage we wished to investigate if this occurs due to its binding to the DNA damage sites. For this, we examined the colocalization of Scm3 with Rad52 upon DNA damage, as the latter is known to be targeted to the damage sites (Lisby et al., 2003; Miyazaki et al., 2004). To visualize both Rad52 and Scm3 simultaneously on chromatin spreads, cells harboring *RAD52-6HA* and *SCM3-13MYC* were mock-treated or treated with 0.02% MMS for 90 mins. We did not observe any significant co-localization between Scm3-13Myc and Rad52-6HA in untreated cells as judged by low Pearson’s Correlation Coefficient value (PCC < 0.4) (Figure 4A-B). However, in the cells treated with MMS, we could observe a significant increase in the PCC value (PCC > 0.5) (Figure 4B). This indicates a partial co-localization between Scm3 and Rad52 in the presence of the DNA damage and implicates that Scm3 may localize to the damage sites in response to DNA damage.

**Figure 4.**
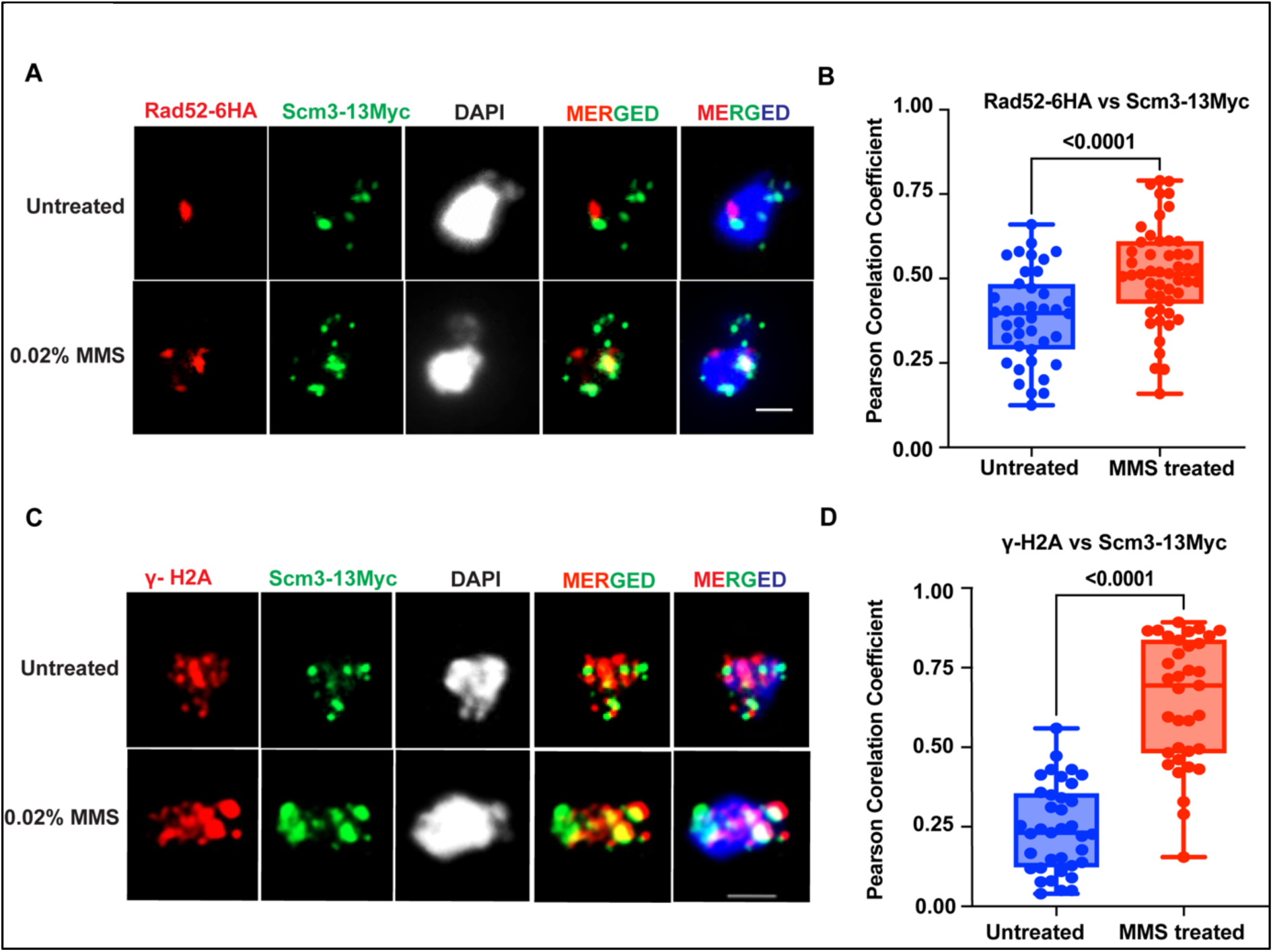
Scm3 colocalizes with DDR proteins in response to DNA damage. **(A)** Representative images showing the localizations of Rad52-6HA and Scm3-13Myc using chromatin spread. The mid-log grown *RAD52-6HA SCM3-13MYC* cells were treated with 0.02% MMS or mock-treated for 90 mins for the assay. **(B)** The quantification of the colocalization between Rad52-6HA and Scm3-13Myc was calculated using Pearson’s correlation coefficient (PCC). **(C)** Representative images showing the localizations of γ-H2A and Scm3-13Myc using chromatin spread. The mid-log grown *SCM3-MYC* cells were treated with 0.02% MMS or mock-treated for 90 mins for the assay. **(D)** The quantification of the colocalization between γ-H2A and Scm3-13Myc was calculated using PCC. Anti-HA (12CA5, Roche), anti-Myc (9E10, Roche), and anti-H2A (ab15083, Abcam) antibodies were used to detect HA, Myc, and H2A-S129, respectively. At least 30 spreads from 2 independent experiments were analyzed for each set in **(B),** and **(D)**. The *p* values were estimated by unpaired two-tailed student’s t-test. Scale bar = 2 µm.

Next, we wished to examine the behavior of Scm3 with respect to another DNA damage marker. In mammals, the DSB sites are marked by γ-H2Ax due to phosphorylation of the H2A variant, H2Ax, by ATM/ATR kinases (Burma et al., 2001; Lee et al., 2014) In yeast that lacks H2Ax, the DSBs are marked by γ-H2A due to phosphorylation of histone H2A by Mec1 and Tel1 kinases (Lee et al., 2014). To be noted, γ-H2A is also found at the heterochromatin region and co-localizes with yeast heterochromatin protein Sir3 (Javaheri et al., 2006; Kirkland et al., 2015; Kitada et al., 2011). We performed a chromatin spread assay using antibodies specific to γ-H2A and observed a drastic increase in the intensity of γ-H2A when the cells were subjected to DNA damage, which is expected (Figure S6 A, B). To verify that the phenomenon is DNA damage associated, we observed a significant increase in the colocalization between Rad52 and γ-H2A in response to DNA damage (Figure S6 C-D). To further correlate the localization of Scm3 with DNA damage, we then visualized both Scm3-13Myc and γ-H2A simultaneously on chromatin spreads from *SCM3-13MYC* cells mock-treated or treated with 0.02% MMS for 90 mins before they were harvested for the assay. We did not observe any significant co-localization between Scm3-13Myc and γ-H2A in untreated cells as judged by low Pearson’s Correlation Coefficient value (PCC < 0.3) but found a significant increase in the value (PCC > 0.6) in MMS treated cells (Figure 4C-D). Taken together, we conclude that the association of Scm3 with chromatin and its co-localization with DDR proteins increases with DNA damage.

### Scm3 is recruited to a site-specific DSB

The overall increased association of Scm3 with the chromatin and its colocalization with Rad52 and γ-H2A in response to MMS-mediated DNA damage prompted us to examine if Scm3 is recruited specifically to DNA damage sites. For this, we utilized the chromatin immunoprecipitation (ChIP) assay to visualize the dynamics of Scm3 recruitment at an induced DSB site. We used a yeast strain (NA14) harboring an HO cut site integrated at the mutant *ura3* locus, which is cleavable by HO endonuclease upon galactose induction (Agmon et al., 2009; Fangaria et al., 2022). The strain also contains a distant *URA3* locus, which is utilized to repair the cut *ura3* locus post-DSB induction. The cleavage by HO endonuclease was first verified by PCR-based analysis using primers complementary to the DNA sequence flanking the HO cut site (Figure 5A). The DNA damage was maximal at 2 hrs of galactose induction, and the damage was majorly repaired by 4 hrs of galactose induction (Figure S7 A, B).

**Figure 5.**
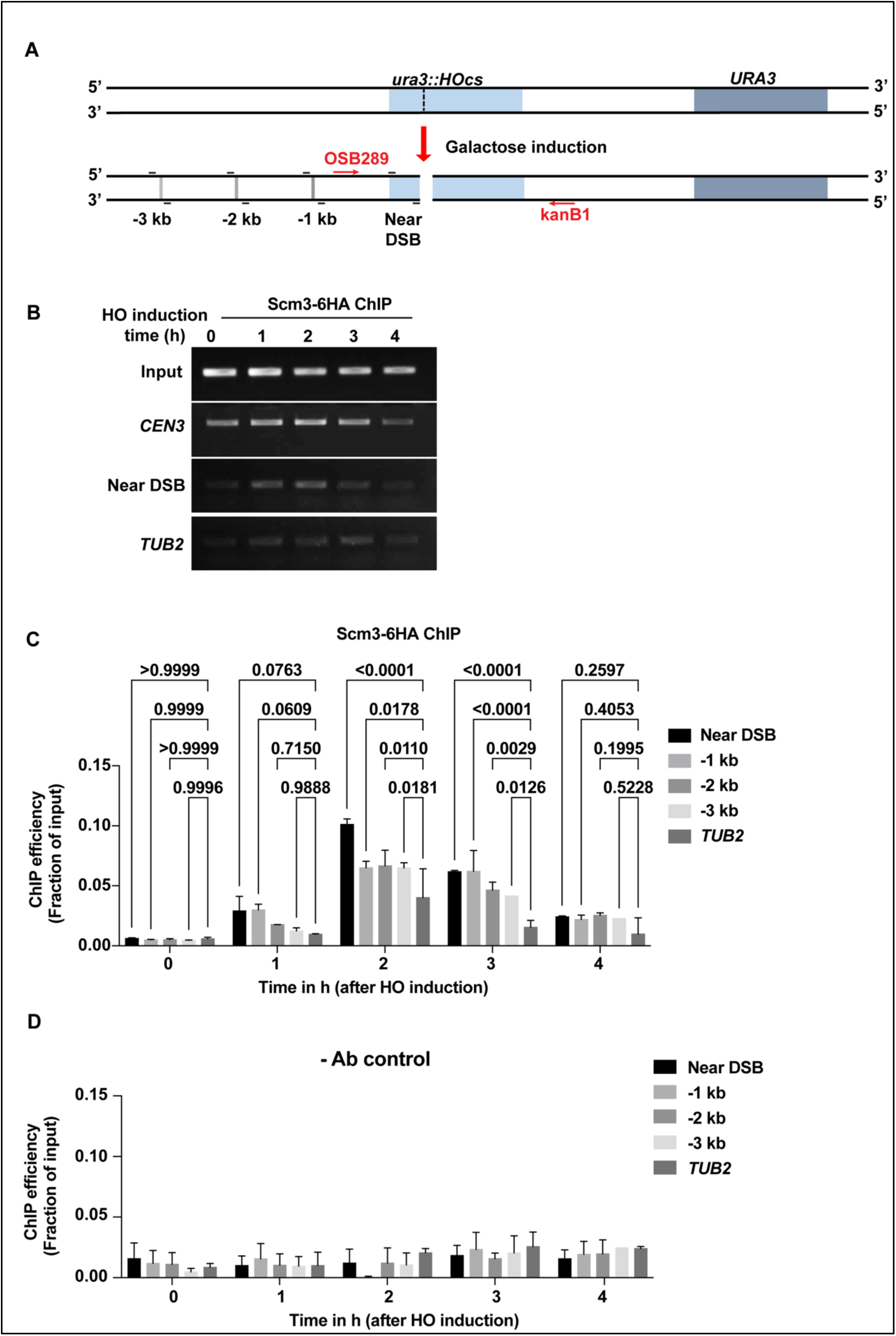
Scm3 is recruited to a site-specific double-stranded break (DSB). **(A)** Schematic representation of a cut site of the *HO* endonuclease (*HOcs*) integrated at the *URA3* locus in the NA14 strain. Upon galactose induction, the *HO* endonuclease cleaves the DNA at the specific site generating DSB. The positions of the primers (OSB289 and kanB1) are shown that were used to detect the DSB, and the binding of Scm3 near and distal (-1 to -3 kb) to the DSB **(B)** ChIP analyses were performed to quantify the association of Scm3-6HA in NA14 strain harboring *SCM3-6HA* with the indicated loci at the indicated time points. The PCR amplified DNA bands following ChIP were run on agarose gel and stained with ethidium bromide. **(C)** ChIP analyses for measuring the association of Scm3-6HA with the indicated loci were performed at the indicated time points of HO induction using anti-HA (12CA5, Roche) antibodies. The ChIP efficiency as percentage enrichment per input obtained by qPCR analysis of the immunoprecipitated DNA was plotted. **(D)** Same analysis as in (C) using no antibody (-Ab) control. The *p* values in **(C)** and **(D)** were estimated from two independent experiments.

We monitored the recruitment of Scm3 near DSB and at sites distal to that up to -3 kb (Figure 5A) by performing ChIP of Scm3-6HA from the samples collected at every hr up to 4 hrs following galactose induction. As a positive control for Scm3 ChIP, we found significant enrichment of Scm3 at the centromere (*CEN3)* (Figure S7C). We observed a significant increase in the Scm3 binding near DSB at 2 hrs of galactose induction and a decrease in the enrichment at 4 hrs, possibly due to the gradual repair of the damage (Figure 5B, C). We also observed an increase in the enrichment of Scm3 up to -3 kb sites distal to the cut site, which decreased gradually with damage repair (Figure 5C) and no significant enrichment at the *TUB2* locus, used as negative control (Figure 5C) or in the no antibody control (Figure 5D). This suggests that Scm3 is maximally recruited to the cleavage sites at the time of DNA damage and gradually dissociates as and when the DNA is repaired. To verify that HO-induced DSB is able to recruit the DNA repair proteins, we also investigated the enrichment of such a protein, Rad51, that is known to bind the DSB sites (Fangaria et al., 2022) and found it is maximally enriched at these sites during 2-3 hrs of post-galactose induction (Figure S7D). We conclude that upon DNA damage Scm3 associates with the damage sites and may act in *cis* to facilitate damage repair.

### Activation of the DDR checkpoint is perturbed in the *scm3* mutant

Having shown the recruitment of Scm3 at the DNA damage sites, we wished to address the role of Scm3 at these sites in case of DNA damage. Since we found that Scm3 depleted cells are compromised in repairing DNA damage, it is possible that being at the damage sites Scm3 activates the DDR checkpoint, which is required for the damage repair. In the DDR pathway, Rad53 is an effector kinase that becomes phosphorylated in a Mec1 sensor kinase-dependent manner in response to DNA damage (Ciccia and Elledge, 2010; Melo and Toczyski, 2002; Zhou and Elledge, 2000). Phosphorylation of Rad53 is essential to halt the cell cycle and provide cells time to repair the damaged DNA. To examine whether Scm3 depleted cells have defects in the activation of DDR checkpoint and in halting the cell cycle, we measured Rad53 phosphorylation status in the wild type and the Scm3 depleted cells after treating them with 0.02% MMS for 90 mins (Figure 6A). Anti-Rad53 antibodies that recognize both phosphorylated and unphosphorylated forms of Rad53 were used. We observed lesser phosphorylation of Rad53 in the *scm3* than in the wild-type cells treated with MMS, but no difference was found in the untreated cells (Figure 6B).

**Figure 6.**
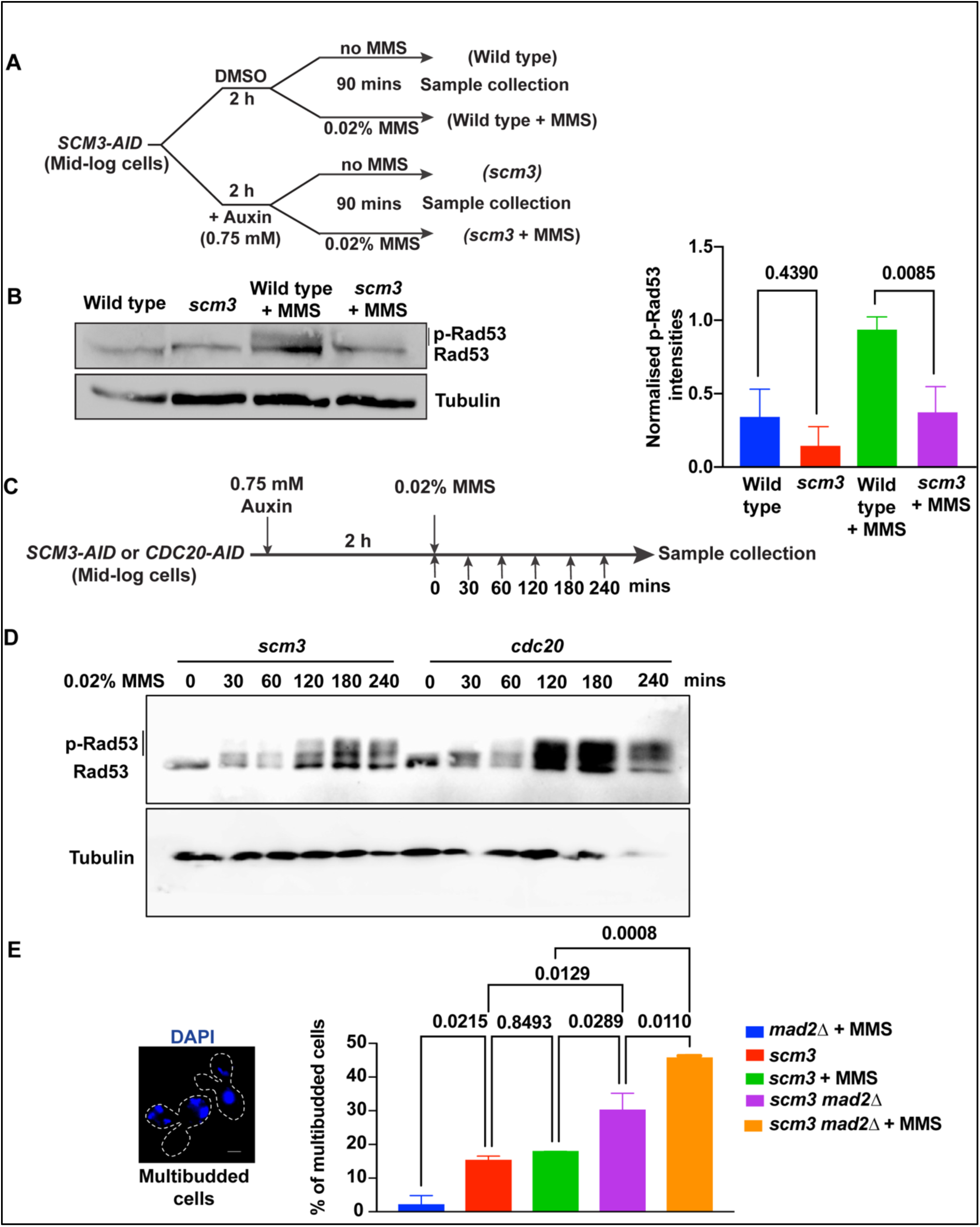
Evidence of improper activation of DNA damage checkpoint in absence of Scm3. **(A)** The schematic of the experimental strategy that was followed for detecting Rad53 in *SCM3-AID* cells in the presence of DMSO (wild-type) or auxin (Scm3 depleted) and each either treated with MMS or left untreated. **(B)** Left, western blots showing p-Rad53/Rad53 and tubulin (as loading control) using anti-Rad53 (ab104232, Abcam) and anti-tubulin (MCA78G, Serotec) antibodies, respectively, from indicated cells obtained from (A). Right, the intensities of the p-Rad53 measured by ImageJ were plotted after normalizing with the corresponding loading control. **(C)** The schematic of the experimental strategy to detect Rad53 in Scm3 or Cdc20 depleted cells treated with MMS for different time points. **(D)** Western blots developed as in (B) from indicated cells obtained from (**C)**. **(E)** *mad2Δ, SCM3-AID,* and *SCM3-AID mad2Δ* cells were first treated with 0.75 mM auxin for 2 hrs to deplete Scm3 and then either treated with 0.02% MMS for 90 mins or left untreated. The cells were harvested and were scored for the presence of multibudded cells (shown on the left). At least 150 cells from three independent experiments were analyzed for each set. The *p* values were estimated by one-way ANOVA test. Scale bar = 2 µm.

Since Scm3-depleted cells, unlike the wild type, arrest at the G2/M stage due to activation of spindle assembly checkpoint (SAC) (Camahort et al., 2007; Stoler et al., 2007), the observed difference in Rad53 phosphorylation could be due to the difference in cell cycle stages. To circumvent this issue, we grew *CDC20-AID* and *SCM3-AID* cells for 3 hrs before treating them with 0.75 mM auxin for 2 hrs to arrest them first at G2/M by depleting Cdc20 and Scm3, respectively (Figure 6C). The cells were then treated with 0.02% MMS and harvested at indicated time points for total protein extraction. The *CDC20-AID* and *SCM3-AID* cultures showed around ∼80% metaphase arrest cells, in 2 hrs of auxin treatment (0 mins in Figure S8A). Notably, while the Cdc20-depleted cells maintained the arrest even with a longer incubation in auxin, the Scm3-depleted cells tend to bypass the arrest after a certain time and showed accumulation of multibudded cells with fragmented DAPI, an arrest-release phenotype (Figure S8B, Figure 6E) Consistent with our previous results (Figure 6C), we observed that in Scm3 depleted cells, Rad53 is less phosphorylated than in Cdc20-depleted cells (Figure 6D), in particular, while comparing the bands at 120 mins when the percentage of metaphase cells was found similar in two cultures (Figure S8B). These results indicate that in the absence of Scm3, when there is DNA damage, the activation of the DDR checkpoint is improper, and consequently, the phosphorylation of Rad53 is compromised.

If the loss of Scm3 fails to activate the damage checkpoint properly, then it is expected that the cell cycle arrest phenotype in response to DNA damage would also be defective and would produce multibudded cells. We wished to examine this possibility. However, Scm3-depleted cells arrest by spindle assembly checkpoint (SAC) at G2/M due to improper kinetochore-microtubule attachment because of failure in Cse4 deposition (Camahort et al., 2007; Stoler et al., 2007). To exclude the effect of SAC-mediated arrest, we deleted the SAC component *MAD2* in *SCM3-AID* cells. These cells were first depleted of Scm3 by treatment with auxin and then, along with *mad2Δ* control cells, were challenged with or without MMS for 90 mins before they were analyzed for percentage of multi budded cells (Figure 6E). While the *mad21* cells in the presence of MMS showed nominal multi budded cells (<5%), the same cells upon depletion of Scm3 showed a steep rise in the phenotype (>40%), suggesting that Scm3 depleted cells fail to activate DNA damage checkpoint and to arrest the cell cycle in the presence of DNA damage.

Since Scm3 depleted cells showed reduced Rad53 phosphorylation, we wished to examine if the lack of Scm3 perturbs Mec1 kinase function that phosphorylates Rad53. Mec1 phosphorylates histone H2A at S129 position upon DNA damage (Lee et al., 2014). However, we failed to observe any significant difference in the phosphorylation status of H2AS129 between Scm3, or Cdc20 depleted cells (Figure S8C), indicating that the Mec1 function is not perturbed in the absence of Scm3.

### Phosphorylation of Scm3 in response to DNA damage

The function of Scm3 in DDR could be mediated by a damage-induced post-translational modification of this protein. To examine if Scm3 becomes modified upon DNA damage, we treated *SCM3-6HA* cells with increasing concentrations of MMS or with 0.02% MMS with increasing duration. We observed slower migrating bands of Scm3 (Scm3*) that became brighter with increasing concentrations of MMS and present very faintly even in the untreated cells (Figure 7A, left panel). We observed an increase in the normalized intensity of Scm3* and Scm3*/Scm3 ratio with increasing MMS concentration (Figure 7A, middle, and right panels). A similar observation was also made when the cells were treated with 0.02% MMS for increasing time (Figure S9A). Since we could detect Scm3* bands in the untreated cells as well, its enrichment in response to MMS could be due to a cell cycle arrest caused by MMS. To understand the dynamics of the appearance of Scm3* bands over different cell cycle stages and in response to MMS, we released the *bar1Δ SCM3-6HA* cells from alpha-factor mediated G1 arrest into the medium in the absence or presence of MMS, as shown in Figure 7B. We harvested the cells at indicated time points and evaluated the cell cycle synchrony using bud morphology and DAPI staining (Figure 7C, Figure S9 B, C), which depicts that the majority of the cells at 0, 30, 60, 90, and at 120 mins are in G1, S, metaphase, metaphase/anaphase and anaphase, respectively. We could faintly observe the Scm3* bands after release from the G1 phase in MMS-free (untreated) media, and the band intensity did not change over the time period. However, when the cells were released in the presence of MMS, we observed an increase in the intensity of the Scm3* bands over time (Figure 7D). These results suggest that the accumulation of Scm3* in the MMS treated cells is due to DNA damage rather than an effect of the pace of the cell cycle. It is possible that Scm3* bands are phosphorylated forms of Scm3 since the protein has several serine/threonine residues. To test this, we challenged the Scm3-6HA cells with MMS, as in Figure 7A, and resolved the proteins on SDS-PAGE with or without Phos-Tag reagent. The presence of Phos-Tag in the gel enables slow mobility of the phosphorylated proteins and thus further differentiates them from their unphosphorylated forms. Figures S9D and E show the migration of the proteins isolated from *SCM3-6HA* cells challenged with different concentrations of MMS in the absence or presence of Phos-Tag reagent, respectively. The mobility of the Scm3* bands was further reduced with respect to the Scm3 band in Phos-Tag gel compared to the normal gel, suggesting that the modification is likely due to phosphorylation. Notably, in several high throughput studies, Scm3 has been shown to be phosphorylated at S39, S64, and S165 residues (Lanz et al., 2021; MacGilvray et al., 2020; Zhou et al., 2021). The S64 phosphorylation of Scm3 was found to be enriched in cells after MMS treatment for 2 hrs as compared to G1 arrested cells, although the authors could find S64 phosphorylation in only 2 out of 4 replicates used in their study (Lanz et al 2021). However, in the absence of further validation in those studies, our observation of a gradual increase in the Scm3* band intensity with increasing dose of MMS convincingly demonstrate DNA damage dependent phosphorylation of Scm3.

**Figure 7.**
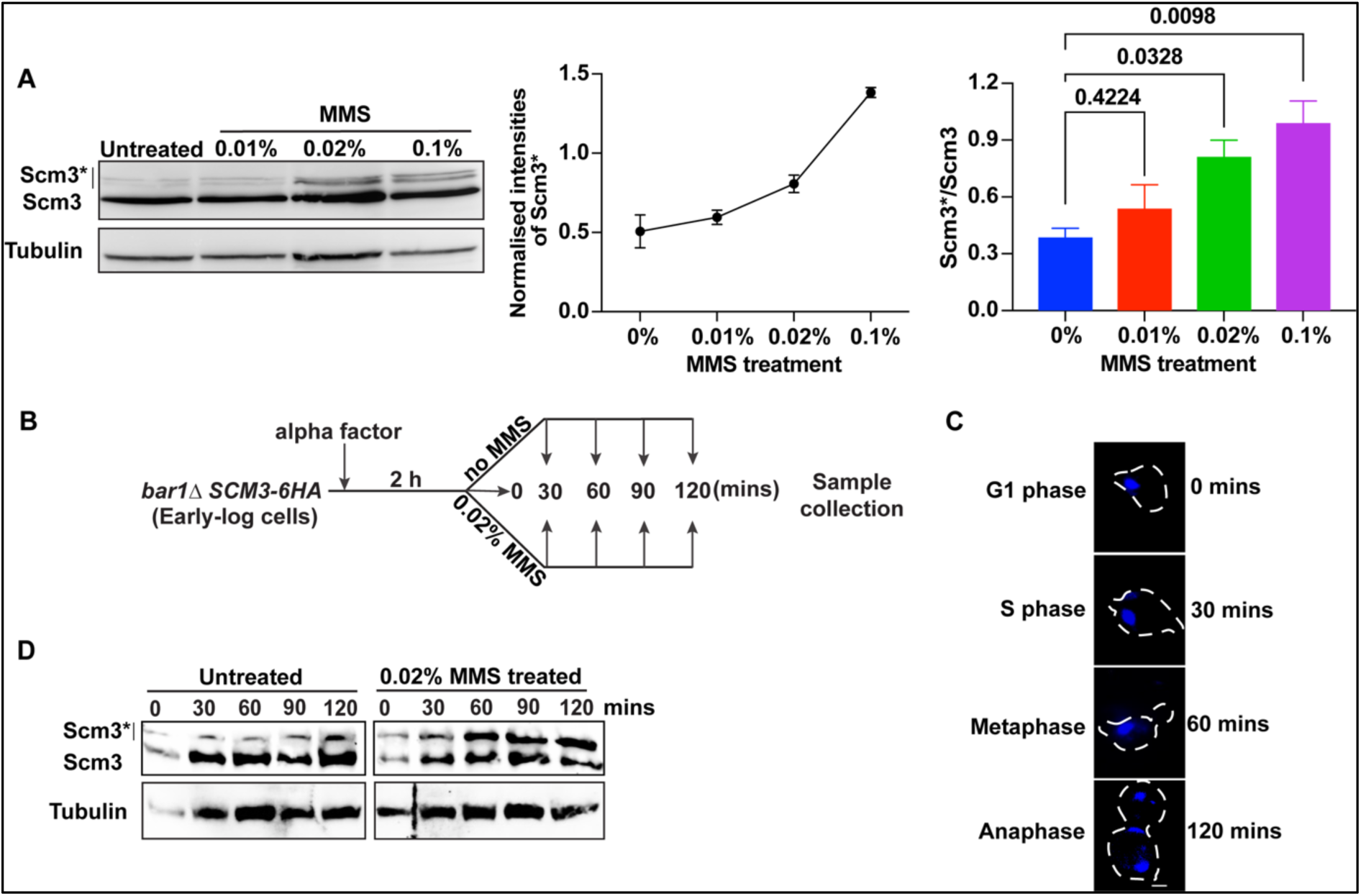
Post-translational modification of Scm3 in response to DNA damage. **(A)** Left, western blots show Scm3-6HA and its modified form (Scm3*) along with tubulin as loading control from the cells untreated or treated with indicated concentrations of MMS for 90 mins. The cell lysates were probed using anti-HA (12CA5, Roche) or anti-tubulin (MCA78G, Serotec) antibodies. Middle, the graph shows the intensity of Scm3* with increasing MMS concentrations, normalized with respect to tubulin using ImageJ software. Right, the graph represents the ratio of the modified to unmodified forms of Scm3. **(B)** The schematic of the experimental strategy that was followed for detecting Scm3-6HA in the G1 arrested cells and following their release from the arrest for the indicated time in the absence or presence of MMS. **(C)** Representative images showing the cell cycle stages by DAPI staining and bud morphology at indicated time points since their release from G1 arrest. **(D)** Western blot showing Scm3-6HA along with tubulin as a loading control from the cells harvested, as shown in (B). The cell lysates were probed using anti-HA and anti-tubulin antibodies to detect Scm3-6HA and tubulin, respectively. Error bars represent the standard deviation from mean values obtained from two independent experiments. Scale bar = 2 µm.

Since, Mec1 kinase majorly contributes to the activation of DDR through protein phosphorylation, we tested if Scm3 undergoes a Mec1-dependent phosphorylation in response to MMS. We constructed the *sml1Δ mec1Δ SCM3-6HA* strain as *mec1Δ* cells are viable in the absence of Sml1 (Zhao et al., 2001, 1998) and challenged this and other control strains with MMS. Briefly, wild type, *SCM3-6HA*, *sml1ΔSCM3-6HA*, and *sml1Δ mec1Δ SCM3-6HA* cells were grown till mid-log and spotted on the plates containing MMS (Figure S9F). As expected, the *sml1Δ mec1Δ SCM3-6HA* cells showed extreme sensitivity to MMS. However, we could not observe any loss of Scm3 phosphorylation in the *sml1Δ mec1Δ* background in the presence of MMS, suggesting that Scm3 phosphorylation in response to DNA damage is not dependent on Mec1 kinase activity (Figure S9G).

## Discussion

The bonafide function of the Cse4 chaperone, Scm3, is to deposit Cse4 at the centromeres during the S phase in budding yeast to facilitate kinetochore formation. The observation that Scm3 persists not only at the centromeres throughout the cell cycle but also associates with the gross chromatin argues for additional kinetochore-independent functions of Scm3 in maintaining genome stability. While in metazoan, Scm3 homolog HJURP was initially identified as a protein involved in DNA damage response (DDR), how HJURP regulates the DDR pathway besides its role in CENP-A deposition is not well understood. The DDR role of Scm3 has never been reported in budding yeast. Whether Scm3 also contributes to DDR pathway in budding yeast is important to address since this organism is distinct from metazoans in terms of harboring small 125 bp point centromeres contrasting megabase pair large regional centromeres in metazoans and having lesser epigenetic control on biological functions, including DNA damage repair. Here, for the first time, we report that Scm3, perhaps upon phosphorylation, promotes DNA damage repair by contributing to the activation of the DNA damage checkpoint, and in the process, it associates with the DDR proteins at the DNA damage sites (Figure 8). Our findings have a significant impact on advancing knowledge of how CENP-A chaperones can be involved in the DDR pathway.

**Figure 8:**
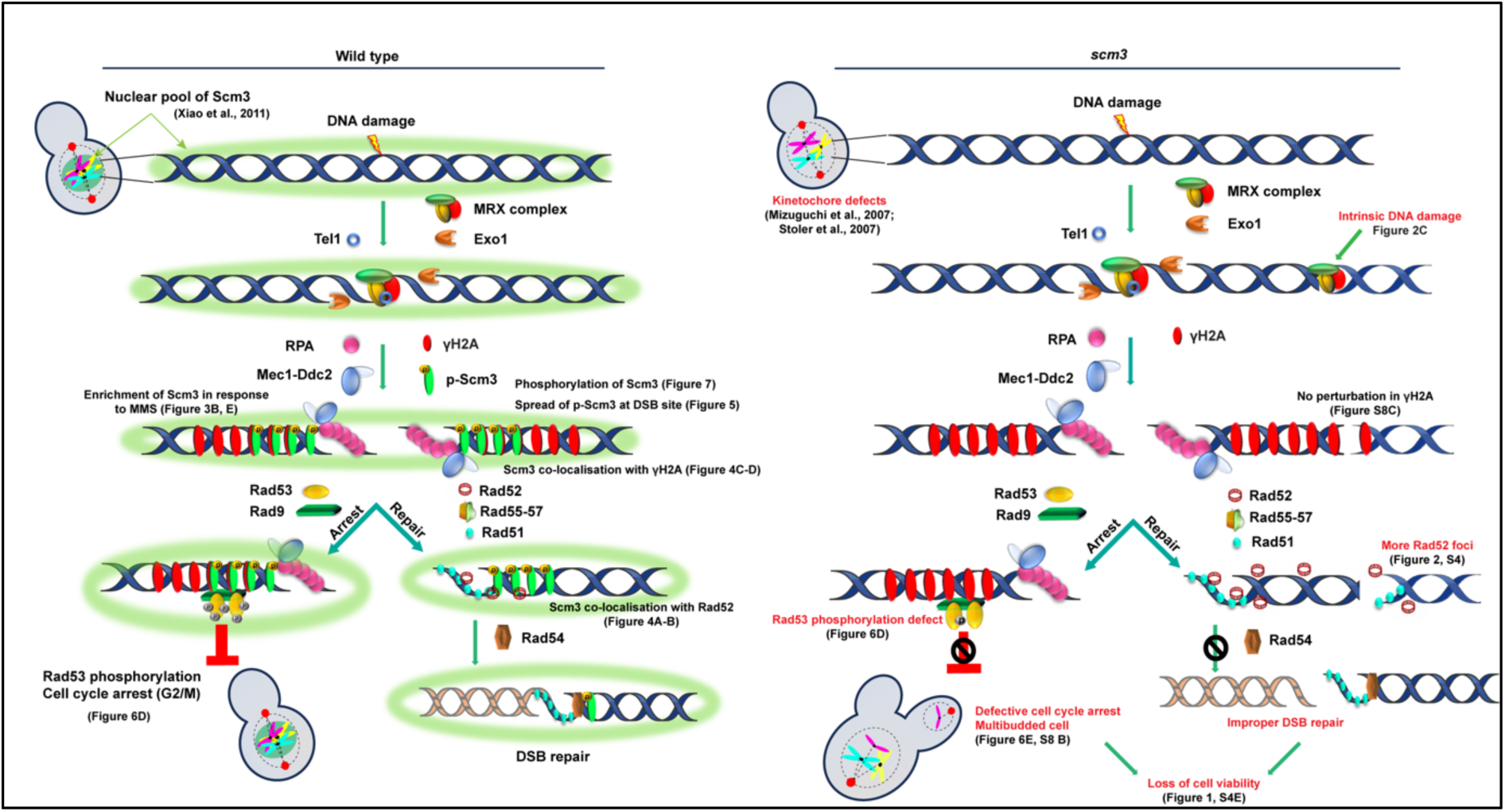
A working model summarizing the role of Scm3 in DNA damage response (DDR). Left, in wild type cells, a subset of the Scm3 molecules present as a nuclear pool (green) are phosphorylated upon DNA damage. The phosphorylated Scm3 (p-Scm3) associates with DSB sites, colocalizes with yH2A, and spreads to the chromatin, flanking the DSB site. Possibly, the p-Scm3 facilitates proper activation of Rad53 through its phosphorylation, leading to cell cycle arrest. Additionally, p-Scm3 co-localizes with Rad52 and assists in DSB repair. Right, in *scm3* mutant cells, there is also an increase in intrinsic DNA damage, corresponding to more Rad52 foci formation, and due to the inability to repair DSBs, the cell viability drops. In parallel, reduced Rad53 activation causes improper DDR-mediated cell cycle arrest generating multibudded cells and aneuploids which causes cell death. The phenotypes associated with the loss of Scm3 are shown in red.

### Lack of Scm3 increases DNA damage sensitivity

It is technically challenging to perform a growth-based sensitivity assay using cells depleted for an essential gene. To circumvent this issue with Scm3, which is essential for cell survival, we used partial depletion of Scm3 to assess the sensitivity of the partially depleted cells to the genotoxic agents, including MMS, HU, and CPT and observed that such cells were sensitive to those agents, (Figure 1, Figure S2A). However, since the primary function of Scm3 is to promote kinetochore formation by depositing Cse4 at the centromeres, it is important to address if the observed sensitivity is due to perturbation in kinetochores. Therefore, we similarly partially depleted two essential kinetochore proteins, Ndc10 and Mif2, and deleted one non-essential kinetochore protein, Ctf19, in separate cells but failed to detect any increased sensitivity to DNA damage stress (Figure S3). These results indicate that the drug sensitivity phenotype of Scm3 depleted cells is not due to weakly formed kinetochores. The arguments favoring the possible centromere-independent DDR functions of Scm3 can be gleaned as follows. From comparing the domain organization of Scm3 with HJURP (Figure S1A), we noticed that besides the CENP-A/Cse4 binding domain, both HJURP and Scm3 have a common DNA binding domain, which is dispensable for CENP-A/Cse4 binding and also for Ndc10 binding in budding yeast. Both in vitro and in vivo evidence are there demonstrating binding of the DNA binding domain to the DNA (Kato et al., 2007; Mizuguchi et al., 2007), suggesting this domain can indeed recruit Scm3 also at the non-centromeric DNA damage sites. In support of this, Scm3 has recently been shown to be phosphorylated in an MMS-dependent manner at S64 (Lanz et al., 2021), which, interestingly, lies within the DNA binding domain. Among other non-essential regions of Scm3, neither the NLS region nor the C-terminal region is involved in DDR (Figure S1B). On the other hand, the D/E-rich region, which is important for Scm3 stability (Stoler et al., 2007), forms a disordered domain as per alpha-fold prediction and hence is believed to be capable of interacting with multiple proteins or can act as a scaffold for engaging several protein-protein interactions which is required in DDR.

MMS halts replication forks due to the formation of DNA adducts, which eventually generate recombinogenic lesions (Conde and San-Segundo, 2008). In yeast, these lesions are repaired primarily through homologous recombination (HR), facilitated by the Rad52 epistasis group of proteins (Pizzul et al., 2022; Sirbu and Cortez, 2013). Rad52, the central component of the HR pathway, deposits Rad51 onto the broken DNA strand, which then searches for the homologous template and undergoes strand exchange to repair the broken DNA (Haber, 2018). In the presence of any DNA damage, Rad52 relocalizes to form bright foci, which mark the repair centres. Strikingly, we observed that the Rad52 foci are increased drastically in the absence of Scm3, even without any external DNA damage (Figure 2C, Left). The maximal accumulation of Rad52-GFP foci was noticed during the 60 mins post-G1 release, which coincides with the timing of DNA replication (Figure S4C), indicating that Scm3 might be required to avoid generating DNA lesions during replication, which itself is genotoxic. In support of this, previously, it was observed that a significant cell population arrests during the S phase in the absence of Scm3 (Camahort et al., 2007). As expected, the accumulation of Rad52-GFP foci was significantly higher in the *scm3* mutant as compared to the wild-type cells when they were treated with MMS, suggesting that cells become more vulnerable to DNA damage in the absence of Scm3 (Figure 2C, Right).

### Association of Scm3 with non-centromeric loci

In addition to the localization of Scm3 at the kinetochores, we also found it at other loci, including at the ribosomal DNA (rDNA) loops (Figure 3A, arrowhead). In budding yeast, chromosome XII, harbors rDNA with around 150 repetitive loci (Dammann et al., 1993; Smith, 2022). Recombination among these repeats excise rDNA circles, and therefore, efficient DNA damage repair through HR at these loci is crucial for preserving genomic stability. Notably, HJURP has been identified in mammals to bind to the rDNA region and maintain its stability (Kato et al., 2007). The degradation of HJURP using siRNA resulted in increased rDNA recombination events and chromosome instability (Kato et al., 2007). The presence of Scm3 at the rDNA loops suggests that Scm3, like HJURP, is also important in regulating the recombination and repair processes at those loci (Figure 3D).

Both Rad51 and Rad52 have been shown to mark the site of recombination and repair (Lee et al., 2003; Lisby et al., 2001; Miyazaki et al., 2004). In our assay, we observed a statistically significant co-localization of Scm3 with Rad52 in response to DNA damage (Figure 4A). Since during DSB repair, multiple lesions are recruited to a few repair centers, which are marked by Rad51 and Rad52, the colocalization of these proteins with Scm3 implicates the recruitment of Scm3 at the repair centers. Moreover, an overall increase in the chromatin association of Scm3 in response to MMS also suggests that Scm3 is recruited to several repair centers (Figure 3B, C, E, S5). However, as expected, no such increase in the chromatin association was observed for an inner kinetochore protein, Ndc10. It is possible that Scm3 might physically interact with Rad51/Rad52 through the HR or DNA binding domain or the D/E rich region present in the protein (Figure S1A). However, we failed to observe any interaction of Scm3 with Rad52 using co-IP assay.

DNA damage sites are epigenetically marked by phosphorylation of H2AS129 (γ-H2A) through the ATM/ATR kinase pathway (Burma et al., 2001; Lee et al., 2014). However, in budding yeast, γ-H2A is also recruited at the heterochromatin regions (Kitada et al., 2011). In unperturbed cells, we could visualize γ-H2A foci (Figure 4C), possibly due to its presence at the stalled replication forks, fragile sites, telomeric regions, or at the heterochromatin regions (Kirkland et al., 2015; Kitada et al., 2011; Rg Bewersdorf et al., 2006). The increase in intensity of γ-H2A upon MMS treatment (Figure S6A, B) indicates the presence of γ-H2A at multiple DNA damage sites. The significantly high co-localization of Scm3 (PCC > 0.6) with γ-H2A in the MMS treated cells, therefore, indicates that Scm3 is indeed recruited to the DNA damage sites (Figure 4D). This was further verified by using an ectopic DSB system, where we showed the recruitment of Scm3 to the induced DSB sites (Figure 5B). Notably, Scm3’s presence was found not only limited to the regions very close to the cut site but also up to 3 kb distal from there. This may be the reason for observing a high co-localization of Scm3 with γ-H2A in response to MMS, as the latter also spreads distal to the cut site (Lee et al., 2014). Although Scm3 binding at the cut site peaked at 2 hrs following HO induction, the time of maximal DNA cleavage, it persisted for at least 3-4 hrs post induction even when ∼60% of the cut DNA was repaired. This suggests that Scm3 is recruited to the sites of DNA damage and persists there during the repair process.

### Scm3 regulates DNA damage checkpoint

We wanted to address the mechanism by which Scm3 might be involved in DDR. The DNA damage checkpoint arrests the cell cycle in response to DNA damage, allowing cells to repair the damage and maintain cell viability. The serine/threonine kinases Mec1/Tel1 in yeast are the sensor kinases that are activated and recruited to the sites of DNA damage. These kinases relay the signal downstream by phosphorylating and activating the effector kinases, including Chk1 and Rad53 (Pizzul et al., 2022; Zhou and Elledge, 2000). The activation of the effector kinases is critical for activating the further downstream substrates that eventually halts the cell cycle. We observed a defect in the Rad53 phosphorylation in response to MMS-mediated DNA damage in the absence of Scm3 (Figure 6B). Failure to activate Rad53 in the presence of damage leads to improper G2/M cell cycle arrest, which can produce the multi-bud phenotype. In the Scm3 depleted cells deleted for Mad2 to remove the ability of SAC mediated arrest (due to kinetochore perturbation), we observed a significant population of multibudded cells (Figure 6E) when they were challenged with MMS. Only *mad2Δ* cells challenged with MMS did not show such multi-bud phenotype, suggesting that Scm3 is important to arrest the cells in response to DNA damage. Next, we wished to know why the Scm3 depleted cells are compromised to phosphorylate Rad53. It is known that Rad9 acts as an adaptor upon phosphorylation by upstream Mec1 kinase to phosphorylate Rad53 (Gilbert et al., 2001; Pellicioli and Foiani, 2005). To test if lack of Scm3 attenuates Mec1 activity, we tested the level of γ-H2A in the Scm3 depleted cells as Mec1 activity is also responsible to generate γ-H2A (Burma et al., 2001; Lowndes and Toh, 2005). However, the level of γ-H2A was not perturbed in the *scm3* mutant, suggesting that Mec1 activity is intact (Figure S8C). Therefore, currently, we are uncertain about the cause of improper Rad53 phosphorylation; a defect in its auto-activation property or in the action of some other mediator kinases, e.g., Tel1, may be responsible, which can be tested. Nevertheless, given the increased phosphorylation of Scm3 in response to DNA damage, it is possible that the p-Scm3 is required to activate Rad53 (Figure 7A). Although we found that Scm3 phosphorylation is Mec1 independent (Figure S9G), it is possible that other kinases are involved. While the involvement of CDKs in the DNA damage checkpoint is not well understood, it is known that CDKs contribute to the phosphorylation of both Rad52 and Rad51 to promote repair (Lim et al., 2020). Given that the centromeric association of Scm3 varies with the cell cycle stage (Mishra et al., 2011), probably due to oscillating levels of CDK activity, the DNA damage-dependent phosphorylation of Scm3 might also be regulated by CDKs. However, this does not rule out the involvement of other kinases including Rad53 and Tel1, which are closely associated with the DNA damage checkpoint pathway. Further investigation is required to precisely determine the kinase involved in the Scm3 phosphorylation.

In summary, we report here that, like HJURP, Scm3 in budding yeast also possesses functions in DDR, indicating that the centromere independent functions of the CenH3 chaperone are evolutionarily conserved. Although the chromatin landscape of budding yeast is far more ‘open’ with lesser involvement of epigenetic determinants than in metazoans, the inherent similarity in the DNA damage repair pathway may necessitate the utilization of these chaperones in a similar fashion in both cell types. Not much has been studied about how HJURP is involved in the DDR pathway, and as we reveal that Scm3 is involved in the proper activation of the DNA damage checkpoint in the presence of DNA lesions (Figure 8), the same can be tested for HJURP in metazoans as abrogation of such surveillance function in humans can lead to aneuploidy and disease states. Our findings on Scm3 will pave the way for future work using yeast as a model to decipher more genome wide functions of the CenH3 chaperone, which is crucial given its genome wide localization and evidence of its modulation in several patients.

## Acknowledgement

We acknowledge Prof. Martin Kupeic from Tel Aviv University, Israel, for providing us with the NA14 strain. SKG is supported by a grant (BT/PR43050/BRB/10/1992/2021) from the Department of Biotechnology (DBT), Govt of India. PA is supported by CSIR fellowship (09/087(0972)/2019-EMR-I).

## Author contributions

PA performed experiments. PA and SKG designed experiments, analyzed and interpreted the data. PA and SKG wrote the manuscript.

## Materials and Methods

### Yeast strains

All *Saccharomyces cerevisiae* strains used in this study are derived from W303 genetic background and are mentioned in Table S1. Standard growth conditions were used to grow and harvest yeast for all the experiments. C-terminal tagging (AID, HA, Myc, GFP) or gene deletions were performed using PCR-based integration at endogenous locations as described before (Janke et al., 2004; Longtine et al., 1998; Wach, 1996). Standard lithium acetate-based yeast transformation methods were employed for all genetic manipulations (Gietz and Woods, 2002).

### Media, reagents and growth conditions

Yeast strains were grown using standard growth conditions (Sherman, 2002). For plates containing MMS, HU and CPT (Sigma Chemicals, Switzerland), the drugs were top-plated on YPD plates at appropriate concentrations. To arrest the cells at the G1 or metaphase stage of the cell cycle, alpha factor (Sigma Chemicals, Switzerland) or nocodazole (Sigma Chemicals, Switzerland) was added to the cells at indicated concentrations. For HO endonuclease induction, strains were grown on a rich medium containing 3% glycerol prior to the addition of 3% galactose. For western blotting, immunoprecipitation, or immunofluorescence studies the primary antibodies used were rat anti-HA (3F10, Roche), mouse anti-HA (12CA5, Roche), mouse anti-Myc (9E10, Roche), rat anti-tubulin (MCA78G, Serotec), rabbit anti-Rad53 (ab104232, Abcam), rabbit anti-Rad51 (PA5-34905, Invitrogen) and rabbit anti-Histone H2A (phospho-S129) (ab15083, Abcam). The secondary antibodies used were Rhodamine (TRITC)-conjugated goat anti-Rat (Jackson), Alexa Fluor 488 conjugated goat anti-Mouse IgG (Jackson), Rhodamine (TRITC)-conjugated goat anti-Rabbit, HRP-conjugated goat anti-mouse IgG (Jackson), HRP-conjugated goat anti-rat (Jackson), HRP-conjugated goat anti-rabbit (Jackson) at appropriate dilutions.

### Cell spotting and viability assay

The cells were grown in standard media conditions till the mid-log phase. The cultures were then 10-fold serially diluted and spotted on the plates containing auxin (dissolved in DMSO) or other DNA-damaging agents. All the reagents were top-plated onto YPD or specific dropout plates. After spotting, the plates were incubated at 30°C for 2-3 days before they were photographed, depending on the growth rate.

For treatment of the cells in liquid culture, mid-log grown cells were treated with auxin (dissolved in DMSO) or MMS for the indicated time points before they were harvested and spotted on YPD plates or plates containing DNA damaging agents. To measure cell viability, an equal number of cells were plated on drug free or drug containing YPD plates. The cells were grown at 30°C for 2 days, and the number of viable colonies was counted. The colonies formed on the drug free plate were taken as 100% viability, and accordingly the viability percentage was calculated and plotted for drug containing plates.

### G1 arrest and release

Cells were arrested in the G1 phase by adding 0.5 μg/mL α-factor to early log phase cells in liquid culture (Amon, 2002). The cells were incubated for 2-3 hours or until at least more than 90% of cells showed shmoo morphology. For the release of the cells from α-factor arrest, the cells were harvested and washed twice with pre-warmed water at 30°C which was followed by washing twice with pre-warmed α-factor free media at 30°C with at least 10 times the volume of the initial culture. The cells were then released in α-factor free media, and samples were collected at different time points.

### Fluorescence imaging

For live cell imaging, the samples were harvested at indicated time points and washed twice in 0.1 M phosphate buffer before imaging. The cells were processed for DAPI staining, as mentioned elsewhere (Mittal et al., 2020; Shah et al., 2023). Typically, the cells were fixed with 4% formaldehyde at RT for 5 mins. The fixed cells were washed twice with 0.1M phosphate buffer (pH 7.5), vortexed in freshly prepared 50% ethanol for 30 seconds, and rewashed with 0.1M phosphate buffer. Before imaging, the cells were resuspended in freshly prepared DAPI (1 μg/ml, Invitrogen) solution for 20 mins. The images were acquired through the Zeiss Axio Observer Z1 fluorescence microscope (63× or 100×) in z-stack mode (0.2-0.5 μm spacing).

### Chromatin spread

Chromatin spread was performed, as mentioned previously (Ma et al., 2023; Mehta et al., 2014; Shah et al., 2023). Briefly, mid-log (O.D._600_ = ∼1.0) grown cells were harvested and washed once with spheroplasting buffer (1.2 M sorbitol in 0.1 M phosphate buffer). The cells were resuspended in 0.5 ml of the same buffer supplemented with 5 µl of β-mercaptoethanol and 12.5 µl of zymolyase 20T (10 mg/ml, MP Biomedicals, USA) and incubated at 30⁰C for 1 h. Once 80-90% of the cells were spheroplasted (by observing under the microscope), the spheroplasting was stopped by adding 1 ml of ice-cold stop solution (0.1 M MES-pH 6.4, 1 mM EDTA, 0.5 mM MgCl_2_, 1 M Sorbitol). The spheroplasts were centrifuged at 2000 rpm for 2 mins and resuspended in 120 µl of ice-cold stop solution. About 60 µl of spheroplasts were mounted on an acid-washed clean glass slide. The cells were treated with freshly made paraformaldehyde solution (4% paraformaldehyde, 3.4% sucrose), followed by 1% Lipsol (LIP Equipment and services) solution for cell lysis. The lysed cells were homogenously spread over the slide and air-dried at RT. After overnight drying, the slides were treated with 0.4% Kodak Photoflow-200 to avoid photobleaching, followed by 5% skim milk as a blocking solution, which was covered with coverslips. The coverslips were removed, and the slides were incubated with primary and secondary antibodies (1:200) for 60 mins with three times PBS washing between antibody treatments. The slides were then incubated with 100 µl of DAPI (1 µg/ml, Invitrogen) for 30 mins. After PBS wash for 5 mins, 100 µl of mounting solution (90% glycerol supplemented with 1 mg/ml p-phenylenediamine) was added over the slides. A clean coverslip was placed over a slide and was sealed with transparent nail paint.

### Microscopic image analysis

The images acquired through the Zeiss Axio (Observer Z1) fluorescence microscope were processed using Zeiss Zen 3.1 software. The z-stacks with the best signal for a particular channel were extracted and merged separately. For intensity calculation, a Region Of Interest (ROI) was drawn around the signal of interest, and the integrated density was determined to get the ‘signal intensity’ for both channels. An ROI of the same size was put elsewhere in the background, and integrated density was determined to get the ‘background intensity.’ Each signal intensity value was normalized with the background intensity values to get the ‘normalized signal intensity’ which was used to plot the graphs (Shah et al., 2023).

For Pearson’s correlation coefficient (PCC) values to determine the extent of co-localization, the merged z-stacks were extracted, and the ‘Coloc’ tool from the ‘Imaris 8.0.2’ software was used. The ‘automatic thresholding option’ in the ‘Coloc’ tool was used to calculate the threshold for each fluorescence emission channel. The PCC values were recorded separately for each individual spread. The colocalization was described as no (PCC <0.1), partial (PCC = 0.3-0.5) and complete (PCC>0.5) (Ma et al., 2023; Prajapati et al., 2018; Zinchuk and Grossenbacher-Zinchuk, 2014).

### Induction of a site-specific DSB

NA14 (Fangaria et al., 2022) cells were grown till 0.3 OD in the presence of 3% glycerol. 5 OD of the cells were harvested (0 h), and the remaining cells were treated with 3% galactose for 4 hrs. 5 OD cells were harvested every hour till 4 hrs. The cells were lysed using glass beads in lysis buffer (2% Triton X-100, 1% SDS, 100 mM NaCl, 10 mM Tris pH 8, and 1 mM EDTA pH 8.0), and total DNA was isolated as described before (Hoffman, 1997) and was solvent extracted using PCI. The genomic DNA was precipitated and finally resuspended in 30 µl 1X TE. The concentration of DNA was measured using a nano spectrophotometer. About 30 ng of DNA was used for PCR to detect the DNA break using primers OSB289 and kanB1 (Fangaria et al., 2022). Primers against *TUB2* ORF were taken as a control.

### ChIP assay and qPCR quantification

ChIP assay was performed as mentioned previously (Fangaria et al., 2022; Kumar et al., 2021; Makrantoni et al., 2019; Mehta et al., 2014). First, cells were grown in the presence of 3% glycerol till 0.3 OD. A total of 60 OD cells were harvested (0 h), and the remaining cells were treated with 3% galactose. A total of 60 OD of cells were harvested every hr till 4 hrs. The cells were then fixed with 1% formaldehyde for 30 mins at 25°C at 100 rpm, followed by quenching by the addition of glycine at a final concentration of 125 mM. The cells were then harvested and washed twice in ice-cold TBS buffer (20 mM Tris–HCl pH 7.5, 150 mM NaCl) and once with 10 ml of FA lysis buffer (50 mM Hepes–KOH pH 7.5, 150 mM NaCl, 1 mM EDTA, 1% Triton X-100, 0.1% Na-deoxycholate) supplemented with 0.1 % SDS. The pellet was resuspended in 500 µl of 1X FA lysis buffer supplemented with 0.5% SDS, 1 mM PMSF, 1X PIC. The cells were lysed using 0.5 mm glass beads using a mini-bead beater (BIOSPEC, India) for 7 cycles (1 min ON, 2 mins OFF on ice). The lysate was collected in a pre-chilled 1.5 ml eppendorf tube and centrifuged twice at 16,000 x g for 15 mins at 4°C. The pellet was resuspended in 300 µl ice-cold 1X FA lysis buffer supplemented with 0.1% SDS, 1 mM PMSF, and 1X PIC. The chromatin was then sheared to 200-600 bps using a water bath sonicator (Diagenode SA, Picoruptor, BC 100,27 LAUDA Germany) for 30 cycles of 30 s ON / 30 s OFF at 4°C. After clarification of the lysate by centrifugation at 16,000 x g for 10 mins at 4°C, 3-5 µg of appropriate antibodies (were added and incubated at 4°C overnight with gentle rotation. Subsequently, 50 µl Protein-A conjugated Sepharose beads were added and incubated for 2 h at 4°C with rotation. The beads were washed twice with IP wash buffer I (1X FA lysis buffer, 0.1%SDS, 275 mM NaCl), followed by one wash in IP wash buffer II (1X FA lysis buffer, 0.1%SDS, 500 mM NaCl, III (10 mM Tris pH 8.0, 250 mM LiCl, 0.5% NP-40, 0.5% sodium deoxycholate, 1 mM EDTA), and 1X TE at room temperature The beads were resuspended in elution buffer and the chromatin was eluted by boiling at 65°C. The eluate was then decrosslinked overnight at 65°C, followed by Proteinase K (SRL, India) treatment for 2 hrs at 45°C. The DNA was purified by PCI-based purification and precipitated overnight in chilled ethanol at -80°C. The enrichment of obtained chromatin fragments was estimated using qPCR (BioRad CFX96, USA) using specific primers targeting specific and non-specific (negative control) chromatin loci, listed in Table S2. As mentioned before (Kumar et al., 2021; Shah et al., 2023) the following equation was used to estimate the percentage of chromatin enrichment. ChIP efficiency = Enrichment/Input X 100; Enrichment/Input = E˄-ΔCT; ΔCT = CT(ChIP) − [CT(Input) − LogEX(D)]; E = primer efficiency value, CT = Threshold values obtained from qPCR, D = Input dilution factor. E was estimated as {[10^(-1/slope)]– 1} from standardization graphs of CT values against dilutions of the input DNA.

### Chromatin immunoprecipitation (ChIP) and sequencing

ChIP assay was performed as mentioned above with certain modifications. The cells were grown till mid-log and treated with nocodazole (20 ug/ml) for 90 mins to arrest cells at metaphase and were subsequently treated with 0.05% MMS for additional 90 mins. The cells were harvested and processed for immunoprecipitation, as mentioned above except Protein-A conjugated dyna beads were used instead of sepharose beads. 10 ng of DNA eluate was taken to prepare the NGS library using a TruSeq ChIP sample preparation kit from Illumina. The library was prepared as per the manufacturer’s protocol. Library QC was performed by measuring the concentration and size of the libraries by using the 4150 Tapestation system from Agilent and Qubit dsDNA HS assay kit (Thermo Fisher Scientific). The libraries were sequenced using the Illumina platform in paired-end mode with a read length of 151 bp by Macrogen Asia Pacific Pte. Ltd, Singapore.

### Bioinformatics analysis

The ChIP-Seq analysis was performed by Genotypic (Bengaluru,India). The raw reads were processed using FastQC (Andrews S., 2010) for quality assessment and pre-processing, which includes removing the adapter sequences and low quality bases (<q30) using TrimGalore (Krueger F., 2015). The pre-processed high quality read data was aligned to reference *Saccharomyces cerevisiae* genome data base using Bowtie, which is an ultrafast, memory-efficient short read aligner geared towards quickly aligning large sets of short DNA sequences (reads) to large genomes. The aligned files (BAM files) were used for peak calling using MACS2 software with input sample as a control. The bedgraph files generated from MACS2 software were used to plot the graphs using the IGV browser (Robinson et al., 2011).

### Protein extraction, gel running and western blotting

Yeast cells harvested at the mentioned time points were washed once with ice-cold water and once with ice-cold 20% TCA. To preserve the phosphorylation moieties, the cells were lysed in the presence of 1X PhosSTOP (Roche). The pellet was resuspended in 200 µl of 20% TCA, and an equal volume of glass beads was added, and cells were lysed using a mini bead beater (BIOSPEC, India) for 7 cycles (1 min ON, 2 mins OFF on ice). The supernatant was collected after puncturing the O-ring tube and centrifugating at 2800 rpm for 2 mins at 4°C. 400 µl of 5% TCA was added to the beads and spun again to collect the supernatant. The total supernatant was spun at maximum speed for 10 mins at 4°C. The precipitate was kept for air-drying for 10-15 mins before resuspending in 2X SDS sample buffer and 1X Tris (pH 8.0) for neutralization. Before loading, the protein was boiled for 5 mins at 95°C and spun at maximum speed for 30 seconds. Standard procedure was followed for SDS-PAGE and western blot transfer.

For the gel containing the PhosTag reagent, 10 mM MnCl_2_ was added to the resolving gel along with 50 µM of the PhosTag reagent (AAL-107, FUJIFILM, JAPAN). Before western blotting, the gel was washed three times with 1 mM EDTA for 10 mins, followed by 2 washes in transfer buffer. The blot was further incubated with primary antibodies at a dilution of 1:2500, while the secondary antibodies were used at 1:5000 dilution.

### Statistical analyses

The data presented in the main figures and the supplementary figures were obtained from two to three independent experiments. The error bars in the individual bar graphs represent the standard error (SE) derived from the standard deviation (SD). The statistical significance (*p*) was determined by two-tailed Student’s t-test, or one-way ANOVA test, as appropriate. When comparing more than one group, a one-way ANOVA test was used; otherwise, Student’s t-test was used. The ‘N’ values denote the total number of cells analyzed from the combined ‘n’ number of replicates of the individual assays. The *p* values less than or equal to 0.05 are categorized as significant differences. The SD, SE, and statistical significance (*p*) values were calculated using automated modules of GraphPad Prism 9.0 (Version 9.4.1) software.

