## Supplementary file 1 for "Evidence of centromeric histone 3 chaperone involved in DNA damage repair pathway"

**Supplementary material**

**Supplementary figures**

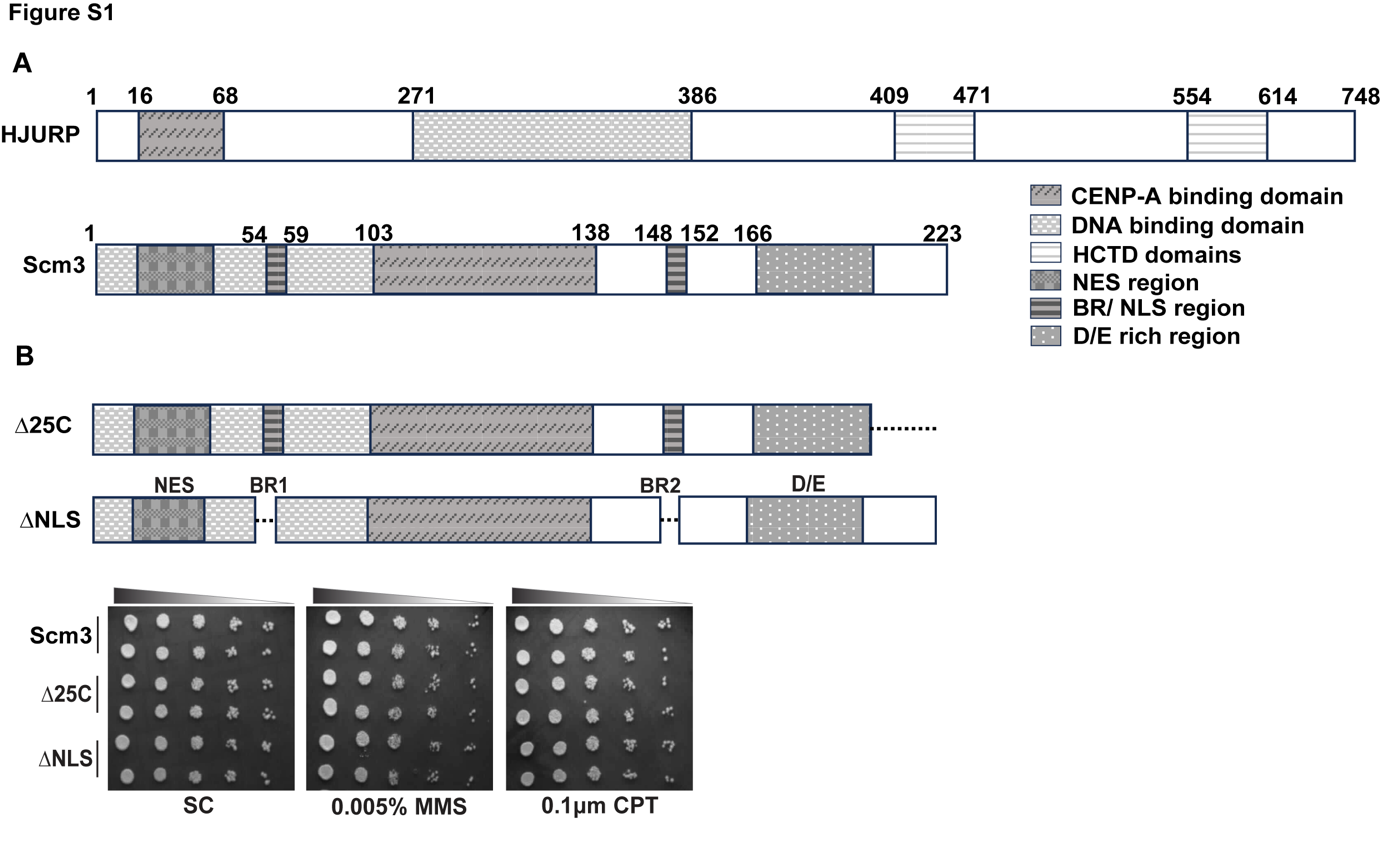

**Figure S1: The NLS domain and the C terminal amino acids of Scm3 are dispensable for the DDR function. (A)** The domain organization of HJURP and Scm3. HCTD, HJURP C-terminal domain); NES, nuclear export signal; BR/NLS, bromodomain or nuclear localization signal; D/E, D/E rich region. **(B)** The domain organization of the Scm3 mutants used for the DNA damage sensitivity assay. Yeast cells harboring indicated Scm3 versions as the sole source of Scm3 were grown till mid-log and were serially diluted and spotted on the indicated plates. The plates were incubated at 30°C for 24-48 hrs before imaging.

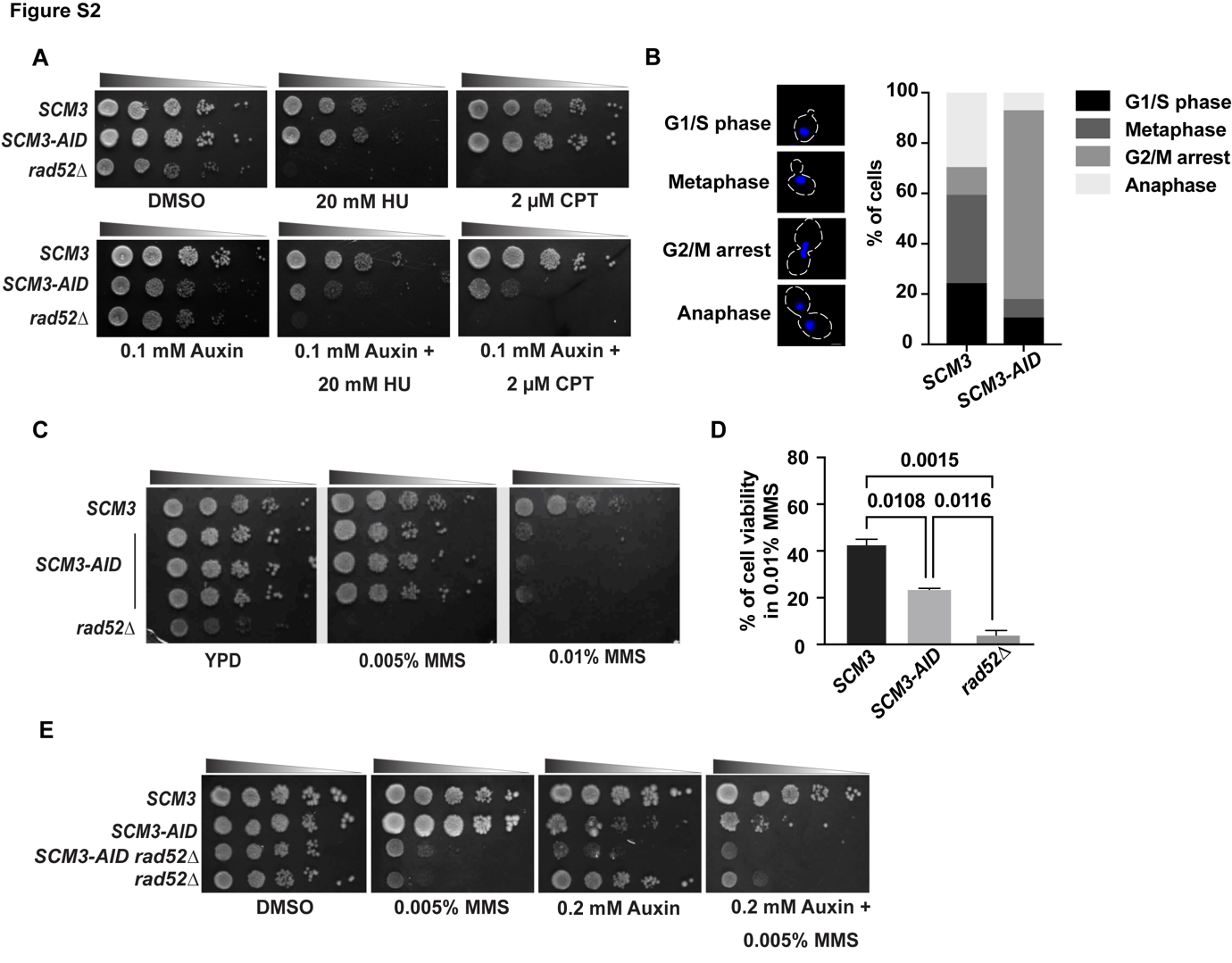

**Figure S2: Cells lacking Scm3 are sensitive to various DNA damaging agents. (A)** *SCM3*, *SCM3-AID, and rad52∆* cells grown till mid-log were serially diluted and spotted on the indicated plates. The plates were incubated at 30°C for 24-48 hrs before imaging. **(B)** *SCM3* and *SCM3-AID* cells were grown till mid log and were treated with auxin for 2 hrs. The cells were harvested and analyzed for DAPI staining and bud morphology to visualize the cell cycle stages. At least 100 cells were analyzed from two independent experiments. **(C)** *SCM3*, *SCM3-AID* (three independent transformants)*, and rad52∆* cells grown till mid-log were pretreated with 0.75 mM auxin for 2 hrs before they were washed, serially diluted, and spotted on the plates supplemented with indicated concentrations of MMS. **(D)** The graph shows the cell viability percentage of the indicated strains treated with 0.01% MMS. An equal number of cells pretreated with 0.75 mM auxin were spread on YPD plates supplemented without or with 0.01% MMS. Taking the number of colonies formed on the YPD plate without MMS as 100% viable condition, the viability on MMS-containing plates was calculated. Error bars were obtained from three independent experiments. The *p* values were estimated by one-way ANOVA test. **(E)** The cells from the indicated strains were grown till mid-log, serially diluted, and spotted on the indicated plates. The plates were incubated at 30°C for 24-48 hrs before imaging.

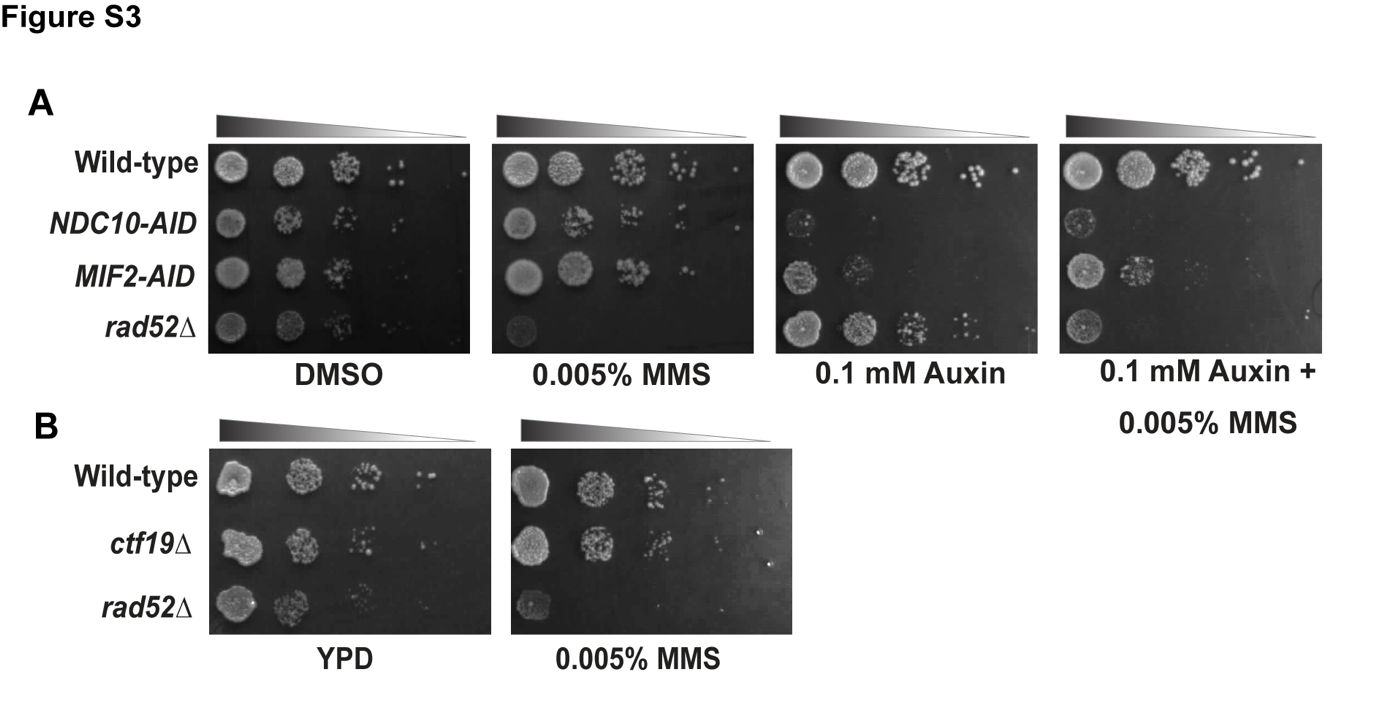

**Figure S3: MMS sensitivity of the cells lacking Scm3 is independent of its kinetochore function. (A)** Wild-type, *NDC10-AID, MIF2-AID, and rad52∆* cells grown till mid-log were serially diluted and spotted on the indicated plates. The plates were incubated at 30°C for 24-48 hrs before imaging. **(B)** Wild-type, *ctf19∆ , and rad52∆* cells grown till mid-log were serially diluted and spotted on the indicated plates. The plates were incubated at 30°C for 24-48 hrs before imaging.

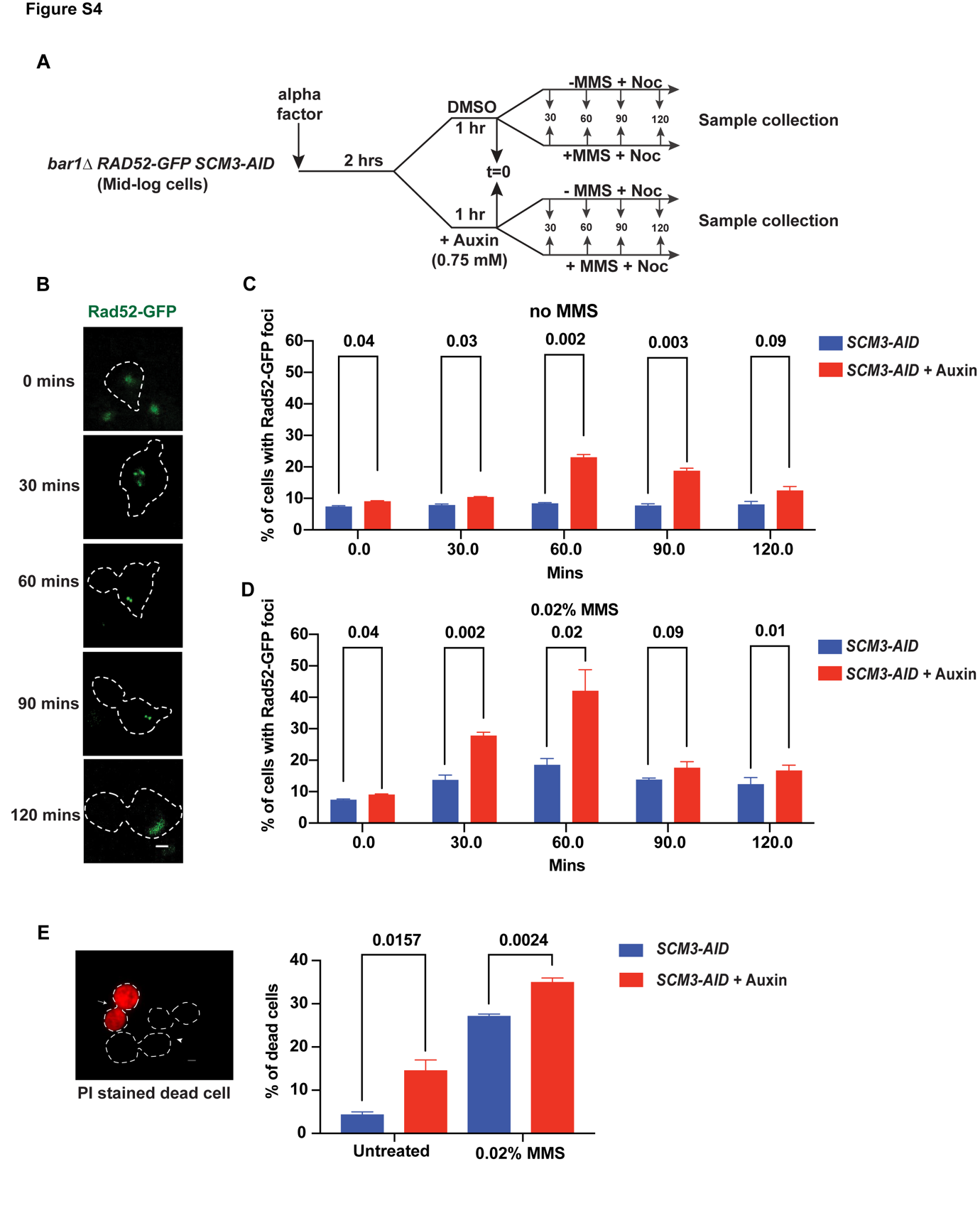

**Figure S4: Loss of Scm3 causes increased Rad52 foci along the cell cycle and cell death. (A)** The schematic of the experimental strategy that was followed for visualization of Rad52-GFP foci in the *bar1∆* *RAD52-GFP SCM3-AID* cells at different stages of the cell cycle in the presence of DMSO (wild-type) or auxin (Scm3 depleted) and each either treated with 0.02% MMS or left untreated. All the cells were arrested eventually at metaphase using nocodazole. **(B)** Representative images showing different patterns of Rad52 foci at indicated time points following their release from alpha factor. **(C, D)** The percentage of Rad52-GFP foci in the indicated cells harvested as shown in (A). **(E)** Cells harvested as shown in Figure 2A were analyzed for cell viability using PI staining. Representative images show PI-positive cells (dead cell, arrow) and PI-negative cells (live cell, arrowhead). The percentage of dead cells with or without MMS treatment for 90 mins are graphically shown. At least 250 cells were analyzed for each set in (C), (D), and (E) from three independent experiments. The *p* values were estimated by unpaired two-tailed student’s t-test. Scale bar = 2 µm.

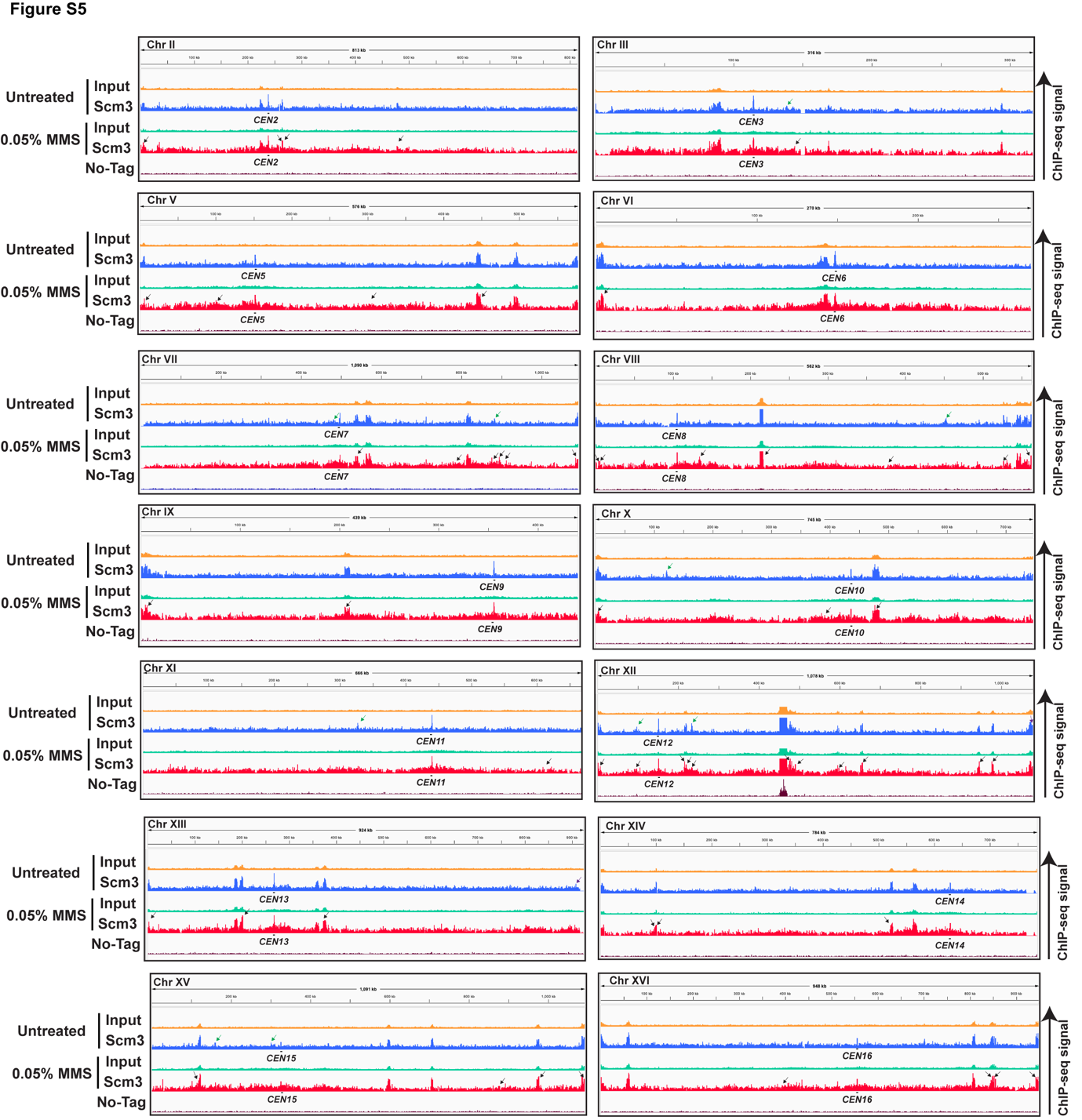

**Figure S5: MMS induced non-centromeric localization of Scm3.** The ChIP-seq signal of Scm3 (Scm3-13Myc) and input from cells either untreated or treated with 0.05% MMS for the indicated chromosomes of yeast; the no-tag strain was used as a control. The green and black arrows represent the non-centromeric localization of Scm3 in untreated and MMS treated samples, respectively. The scale is 7–50 for the input samples and 7-25 for IP samples.

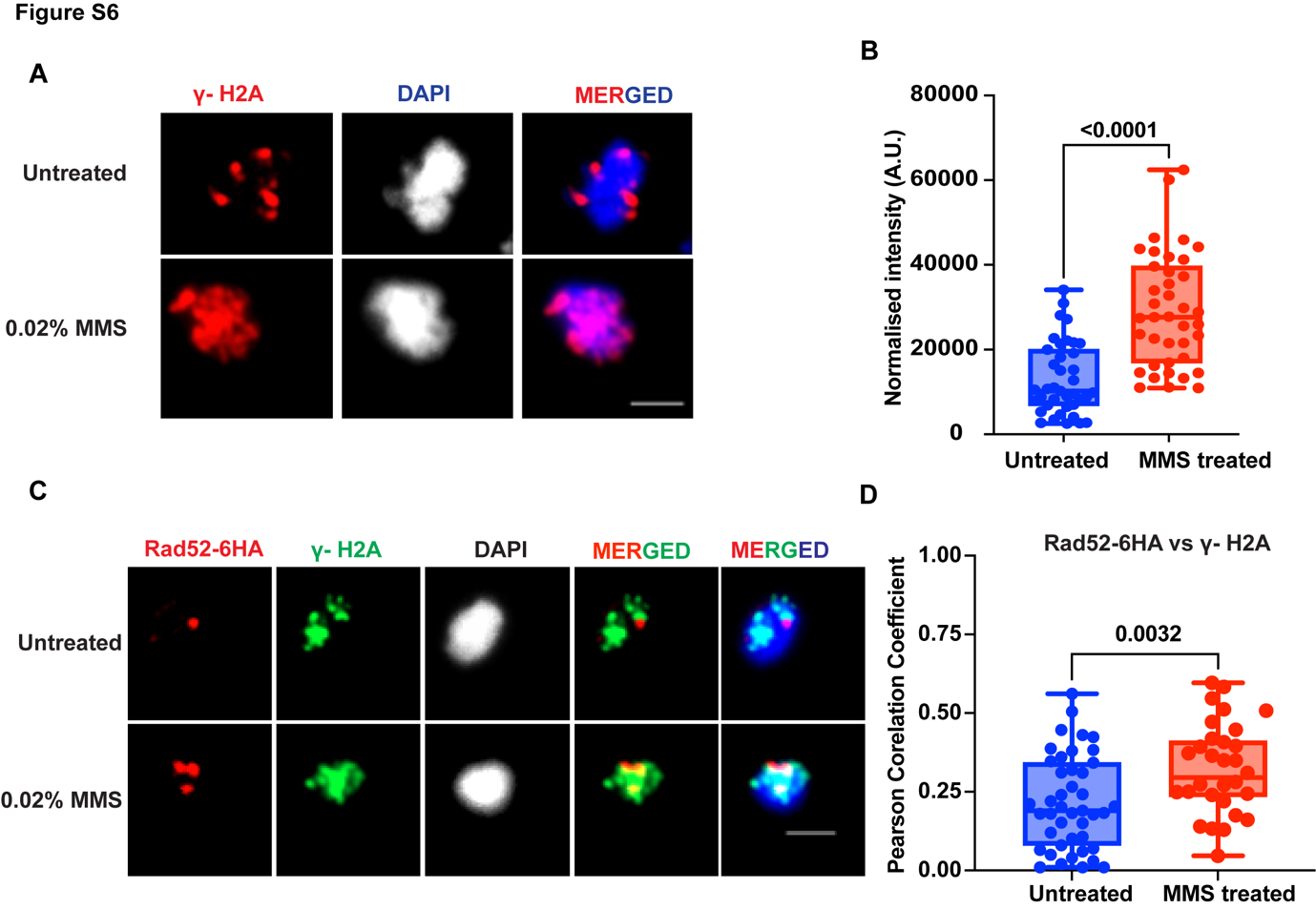

**Figure S6: Yeast γ-H2A marks the sites of DNA damage. (A)** Representative images showing the localization of γ-H2A using chromatin spread. The mid-log grown wild type cells were treated with 0.02% MMS or mock-treated for 90 mins for the assay. **(B)** The intensity of γ-H2A normalized with the background in the presence or absence of MMS was quantified using ImageJ software. **(C)** Representative images showing the localizations of γ-H2A and Rad52-6HA using chromatin spread. The mid-log grown *RAD52-6HA* cells were treated with 0.02% MMS or mock-treated for 90 mins for the assay. **(D)** The quantification of the colocalization between γ-H2A and Rad52-6HA was estimated using Pearson's correlation coefficient (PCC). At least 30 spreads were analyzed for each set in (B) and (D) from three independent experiments. The *p* values were estimated by unpaired two-tailed student’s t-test. Scale bar = 2 µm.

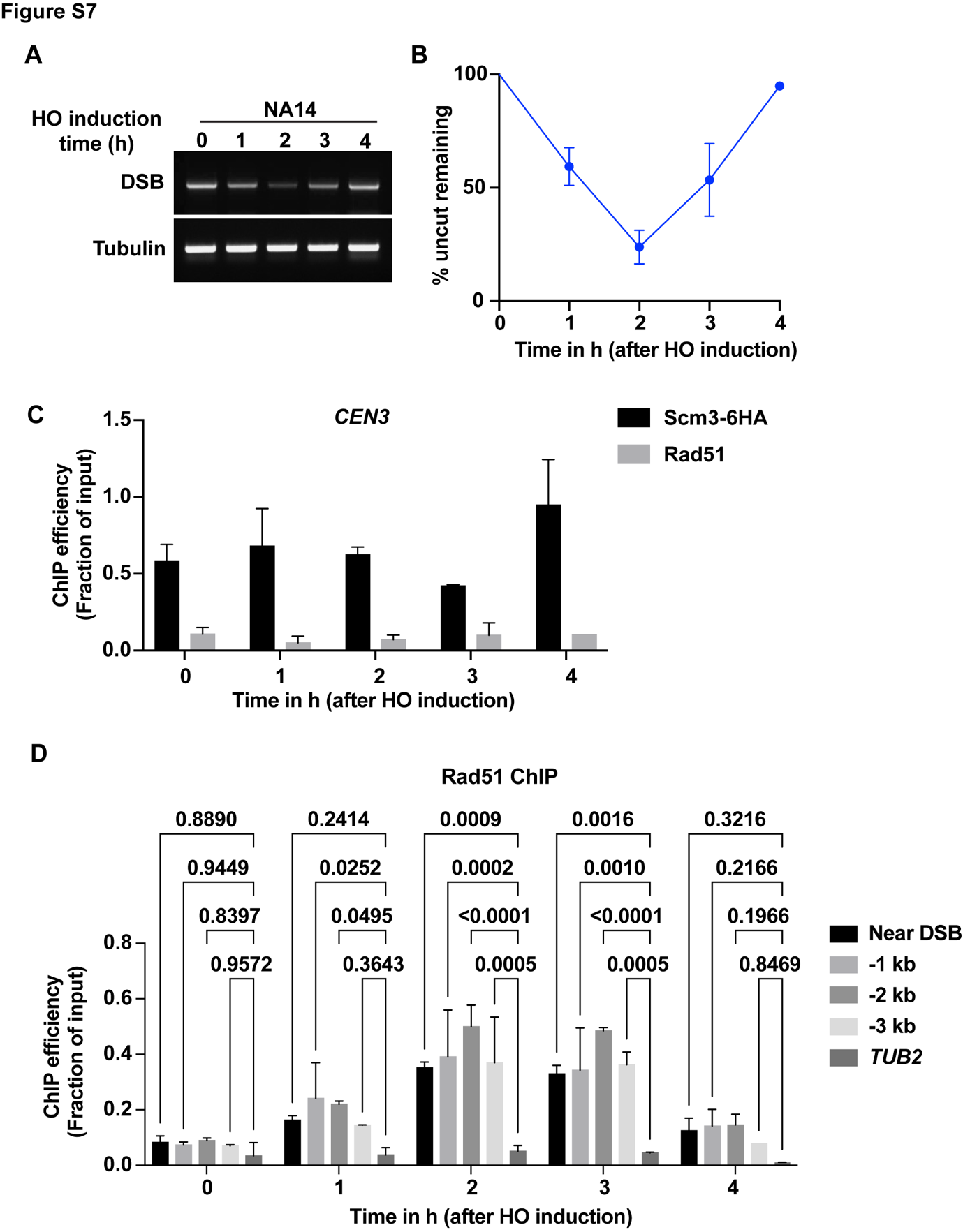

**Figure S7: Dynamics of *HO*-induced DSB formation and protein binding at centromere and DSB sites. (A)**. NA14 yeast cells were grown till mid-log in 3% glycerol, following which galactose was added for *HO* induction. Cells were harvested at indicated time points, and genomic DNA was extracted. PCR was performed to get the band (DSB) to estimate the extent of DNA damage using the primers OSB289 and kanB1, as shown in Figure 4A**.** PCR was also performed using primers against *TUB2* ORF to use as a loading control band (Tubulin). **(B)** The intensity of the DSB band normalized with the tubulin band in (A) was quantified using ImageJ software. The value at 0 hr was taken as 100% DSB band remaining (uncut), and accordingly, the % uncut DNA remaining for other time points was calculated. **(C)** ChIP analyses for measuring the association of Scm3-6HA and Rad51 with the *CEN3* locus were performed at the indicated time points of HO induction using anti-HA (12CA5, Roche) and anti-Rad51 (PA5-34905, Invitrogen) antibodies. **(D)** ChIP analyses for measuring the association of Rad51 with the indicated loci were performed at the indicated time points of HO induction using anti-Rad51 antibodies. The error bars in B, C, and D are estimated from two independent experiments.

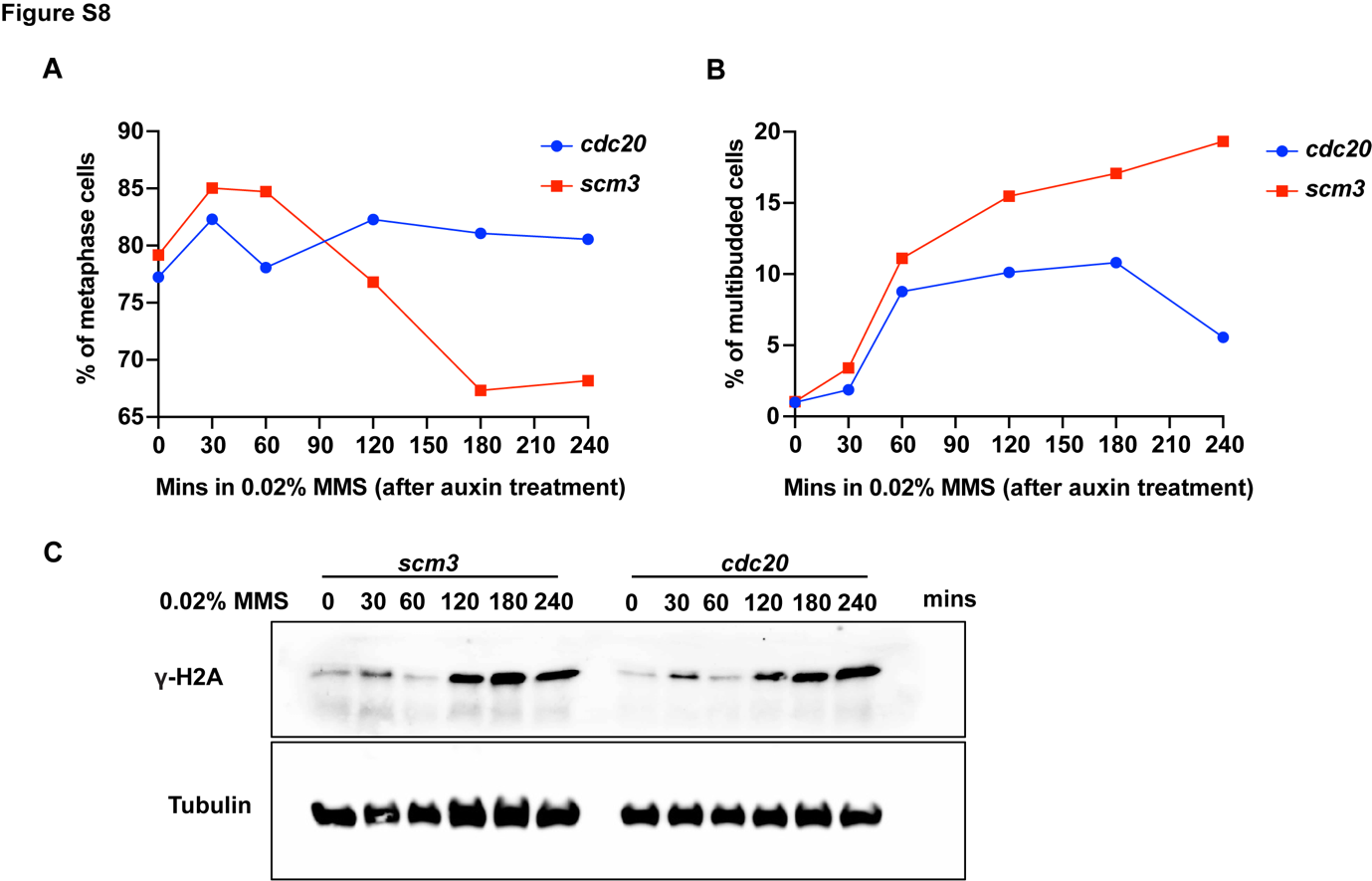

**Figure S8: H2A phosphorylation is not perturbed in the absence of Scm3. (A)** *CDC20-AID* or *SCM3-AID* strains were grown till mid-log and treated with 0.75 mM auxin for depletion of Cdc20 or Scm3, respectively. The cells were then treated with MMS for 240 mins, and samples were harvested at indicated time points. The percentage of metaphase cells judged by DAPI staining and bud morphology for each time point is presented graphically. **(B)** The percentage of multibudded cells in the samples harvested in (A) is presented. **(C)** Western blots showing y-H2A and tubulin (as loading control) using anti-p-H2AS129 (ab15083, Abcam) and anti-tubulin (MCA78G, Serotec) antibodies, respectively. Cells were harvested following strategy given in Figure 5C. At least 100 cells from two independent experiments were analyzed for each set in **(A, B).**

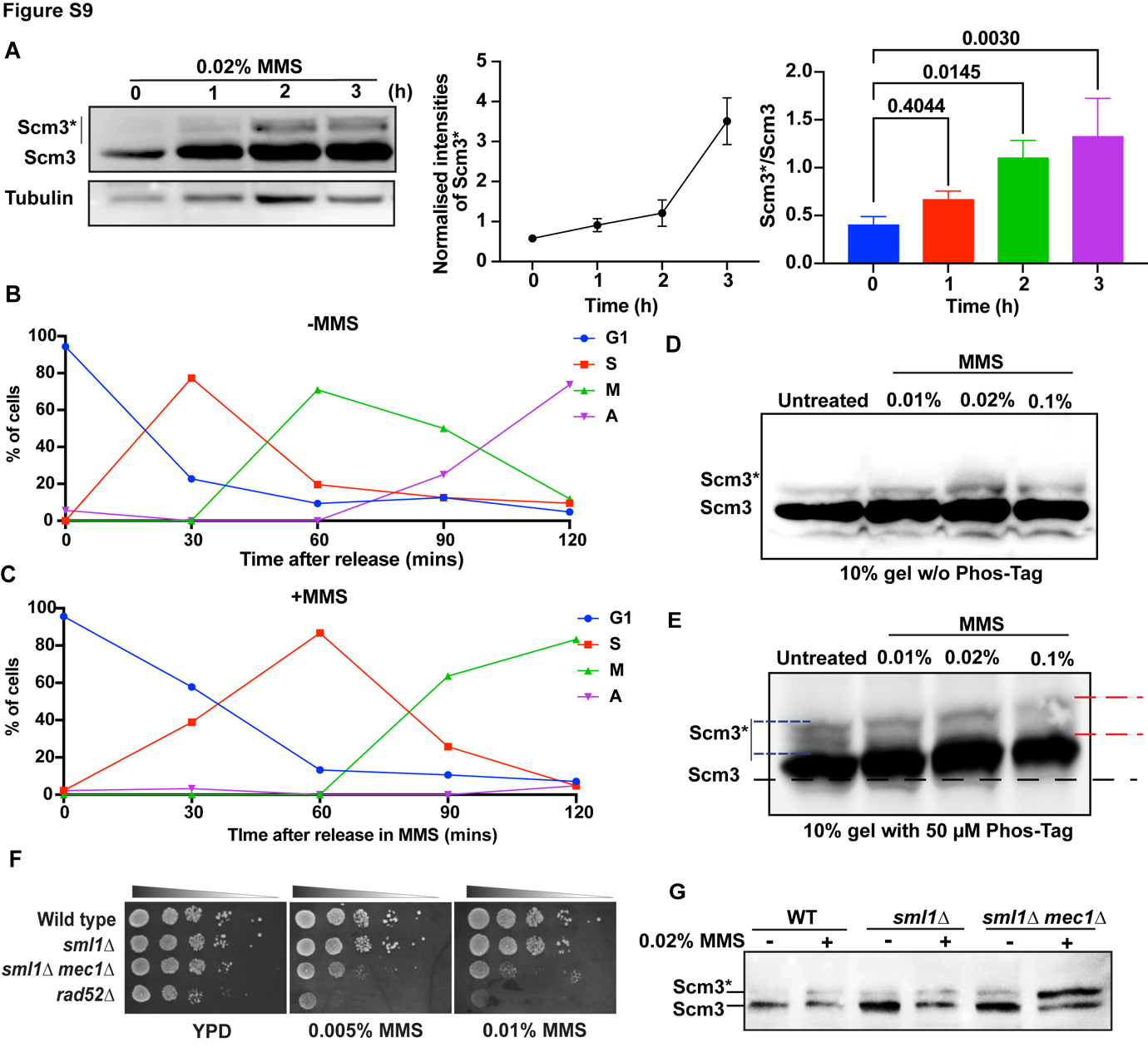

**Figure S9: Phosphorylation of Scm3 in response to DNA damage. (A)** Left, western blots showing Scm3-6HA and its modified form (Scm3*) along with tubulin as loading control from *SCM3-6HA* cells untreated or treated with 0.02% MMS for indicated time points. The cell lysates were probed using anti-HA (12CA5, Roche) or anti-tubulin (MCA78G, Serotec) antibodies. Middle, the graph showing the intensity of Scm3* with increasing duration of MMS treatment, normalized with respect to tubulin using ImageJ software. Right, the graph representing the ratio of the modified to unmodified forms of Scm3. **(B, C)** *bar1∆ SCM3-HA* cells were released from G1 arrest by alpha factor into medium in absence (B) or presence (C) of MMS. The cell cycle stages of the cells at the indicated time points were determined using DAPI staining and bud morphology and are graphically shown. **(D, E)** *SCM3-6HA* cells were treated with the indicated concentration of MMS for 90 mins and total protein was extracted. The proteins were run on 10% SDS-PAGE without (D) or supplemented with 50 µm Phos-Tag reagent (E). The Scm3 band under untreated condition (black dotted line) gradually shifts upwards with increasing concentration of MMS in the presence of Phos-tag reagent, shown by a basal black line. Two red dotted lines show an increase in the spread of Scm3* in 0.1% MMS lane as compared to the same shown by two blue lines in untreated lane. **(F)** Mid-log grown wild-type, *sml1∆, mec1∆ sml1∆, rad52∆* cells harboring *SCM3-6HA* were serially diluted and spotted on the indicated plates which were incubated at 30°C for 24-48 hrs before imaging. **(G)** Western blots showing Scm3-6HA and its modified form (Scm3*) from the cells used in (F)**,** untreated or treated with 0.02% MMS for 90 mins.

**Supplementary tables**

**Table S1: List of strains used in this study:**

| **S.no.** | Name | **Genotype** | Source |
| --- | --- | --- | --- |
| **1.** | CRY1 | *MATa, ura3-1 leu2,3-112 his3-1 trp1-1 ade2-1 can1-100* | (Rothstein, 1983) |
| **2.** | SGY10016 | *MATa, ura3-1 leu2,3-112 his3-1 trp1-1 ade2-1 can1-100 rad52∆::KanMx* | This study |
| **3.** | SGY10035 | *MATa, ura3-1 leu2,3-112 his3-1 trp1-1 ade2-1 can1-100 SCM3-AID::KanMx, pADH-Os-TIR1::URA3* | This study |
| **4.** | SGY10037 | *MATa, ura3-1 leu2,3-112 his3-1 trp1-1 ade2-1 can1-100 SCM3-AID::KanMx,pADH-Os-TIR1::URA3, rad52∆::HPHMX* | This study |
| **5.** | SGY10157 | *MATa, ura3-1 leu2,3-112 his3-1 trp1-1 ade2-1 can1-100 ctf19∆::HPHMX* | This study |
| **6.** | SGY71 | *MATa, ho::Lys2, lys2, ura3, leu2::hisG, his3::hisG, trp1::hisG, MIF2-AID::KanMx, pADH-Os-TIR1::URA* | (Mehta et al., 2014) |
| **7.** | SGY79 | *MATa, ho::Lys2, lys2, ura3, leu2::hisG, his3::hisG, trp1::hisG, NDC10-AID::KanMx, pADH-Os-TIR1::URA* | (Mehta et al., 2014) |
| **8.** | SGY10044 | *MATa, ura3-1 leu2,3-112 his3-1 trp1-1 ade2-1 can1-100 SCM3-AID::KanMx, pADH-Os-TIR1::URA, RAD52-GFP::TRP1* | This study |
| **9.** | SGY10061 | *MATa, ura3-1 leu2,3-112 his3-1 trp1-1 ade2-1 can1-100 SCM3-AID::KanMx, pADH-Os-TIR1::URA, RAD52-GFP::TRP1, bar1∆::LEU2* | This study |
| **10.** | SGY10065 | *MATa, ura3-1 leu2,3-112 his3-1 trp1-1 ade2-1 can1-100 NDC10-6HA::HIS3, SCM3-13MYC::KanMx* | This study |
| **11.** | SGY10063 | *MATa, ura3-1 leu2,3-112 his3-1 trp1-1 ade2-1 can1-100 RAD52-6HA::HIS3, SCM3-13MYC::KanMx* | This study |
| **12.** | NA14 | *MATa-inc ura3-HOcs lys2::ura3-HOcs-inc ade2-1 ade3:: GALHO leu2-3112 his3-11,15 trp1-1 can1-100* | (Agmon et al., 2009; Fangaria et al., 2022) |
| **13.** | SGY10179 | *NA14, ura3-1 leu2,3-112 his3-1 trp1-1 ade2-1 can1-100 SCM3-6HA::HPHMX* | This study |
| **14.** | SGY10032 | *MATa, ura3-1 leu2,3-112 his3-1 trp1-1 ade2-1 can1-100 CDC20-AID::KanMx, pADH-Os-TIR1::URA3* | This study |
| **15.** | SGY10038 | *MATa, ura3-1 leu2,3-112 his3-1 trp1-1 ade2-1 can1-100 SCM3-AID::KanMx, pADH-Os-TIR1::URA3, mad2∆::HPHMX* | This study |
| **16.** | SGY10019 | *MATa, ura3-1 leu2,3-112 his3-1 trp1-1 ade2-1 can1-100 SCM3-6HA::HIS3* | This study |
| **17.** | SGY10059 | *MATa, ura3-1 leu2,3-112 his3-1 trp1-1 ade2-1 can1-100 SCM3-6HA::HIS3 bar1∆::LEU2* | This study |
| **18.** | SGY10088 | *MATa, ura3-1 leu2,3-112 his3-1 trp1-1 ade2-1 can1-100 SCM3-6HA::HIS3 sml1∆::LEU2* | This study |
| **19.** | SGY10089 | *MATa, ura3-1 leu2,3-112 his3-1 trp1-1 ade2-1 can1-100 SCM3-6HA::HIS3 sml1∆::LEU2 mec1∆::KanMx* | This study |
| **20.** | SGY10119 | *MATa, ura3-1 leu2,3-112 his3-1 trp1-1 ade2-1 can1-100 Δbar1 scm3Δ::TRP1 pRS423-SCM3::HIS3* | This study |
| **21.** | SGY10120 | *MATa ura3-1 leu2,3-112 his3-1 trp1-1 ade2-1 can1-100 Δbar1 scm3Δ::TRP1 pRS423-scm3-Δ25C::HIS3* | This study |
| **22.** | SGY10121 | *MATa ura3-1 leu2,3-112 his3-1 trp1-1 ade2-1 can1-100 Δbar1 scm3Δ::TRP1 pRS423-scm3-ΔNLS::HIS3* | This study |

**Table S2: List of primers used in ChIP-qPCR:**

| **Name** | **Description** | **Primer sequence (5′-3′)** | **Source** |
| --- | --- | --- | --- |
| *CEN3* | Forward | GATCAGCGCCAAACAATATGG | (Mehta et al., 2014) |
|  | Reverse | AACTTCCACCAGTAAACGTTT |  |
| *CEN4* | Forward | GCTTGCAAAAGGTCACATGC | (Mehta et al., 2014) |
|  | Reverse | GAGCAGGTTTTATGTTTCGG |  |
| *TUB2* | Forward | CTTGTAGACAGCGTCATGG | (Mehta et al., 2014) |
|  | Reverse | CAGATGTCATAAAGTGCTTCG |  |
| DSB | Forward (OSB289) | GTTAGTTGAAGCATTAGGTCC | (Fangaria et al., 2022) |
|  | Reverse (kanB1) | TGTACGGGCGACAGTCACAT |  |
| Near DSB | Forward | ATGTCGAAAGCTACATATAAG | (Fangaria et al., 2022) |
|  | Reverse | AATGCTTCAACTAACTCCAG |  |
| -1 kb | Forward | GGAGAATCCATACAAGAAATCG | (Fangaria et al., 2022) |
|  | Reverse | CATCTCATTAGTTGGAATTTCG |  |
| -2 kb | Forward | TTTGGTAGATCATTTAAGGGTC | (Fangaria et al., 2022) |
|  | Reverse | CAGGAGATGGCTTAGGCAAG |  |
| -3 kb | Forward | TTTCATTGCTTCCGACTCCG | (Fangaria et al., 2022) |
|  | Reverse | TACGCAGACGAGAAGGCTTC |  |
